# Ensemble docking for intrinsically disordered proteins

**DOI:** 10.1101/2025.01.23.634614

**Authors:** Anjali Dhar, Thomas R. Sisk, Paul Robustelli

## Abstract

Intrinsically disordered proteins (IDPs) are implicated in many human diseases and are increasingly being pursued as drug targets. Conventional structure-based drug design methods that rely on well-defined binding sites are however, largely unsuitable for IDPs. Here, we present computationally efficient ensemble docking approaches to predict the relative affinities of small molecules to IDPs and characterize their dynamic, heterogenous binding mechanisms at atomic resolution. We demonstrate that these ensemble docking protocols accurately predict the relative binding affinities of small molecule α-synuclein ligands measured by NMR spectroscopy and generate conformational ensembles of ligand binding modes in remarkable agreement with experimentally validated long-timescale molecular dynamics simulations. Our results display the potential of ensemble docking approaches for predicting small molecule binding to IDPs and suggest that these methods may be valuable tools for IDP drug discovery campaigns.

## Introduction

Protein structure determination is often the first step in rational drug design. In conventional drug discovery campaigns, ligands are designed to complement the three-dimensional (3D) structures of well-defined binding pockets (1–3). Intrinsically disordered proteins (IDPs), however, do not adopt stable tertiary structures under physiological condition. Instead, they exist as heterogenous ensembles of rapidly interconverting conformations (4–9). IDPs, which make up ∼30% of the human proteome, have essential cellular functions in transcriptional regulation and cellular signaling and mediate the formation of biomolecular condensates (6–8). IDPs are implicated in numerous human diseases and represent a large pool of drug targets that are currently inaccessible to conventional structure-based drug design approaches.

Several small molecules have been discovered that bind IDPs and inhibit their interactions (10–18), and several small molecule IDP ligands have entered human trials (19–21). A preponderance of biophysical evidence suggests that these ligands do not induce their targets to fold into structured conformations upon binding and that these IDPs remain disordered while interacting with small molecule inhibitors discovered thus far (10–16). This has spurred the development of new molecular recognition paradigms, where the affinity of IDP ligands is conferred through dynamic networks of transient interactions that only subtly shift the conformational ensemble of IDPs (10–14, 22–25). This suggests that it may not be possible to identify a small number of representative IDP ligand binding modes to use as starting points for conventional structure-based drug design in IDP drug discovery efforts.

All-atom molecular dynamics (MD) computer simulations, validated by experimental data from nuclear resonance (NMR) spectroscopy, have provided valuable atomic-resolution descriptions of IDP ligand binding modes (11–14, 22–25) and have been found to correctly predict the relative binding affinities of small molecule ligands to some IDP targets (11–13, 24). All-atom MD simulations of IDP ligand binding are, however, computationally expensive. The computational cost of all-atom MD simulations makes this technique relatively impractical for screening large ligand libraries to discover novel inhibitors. Molecular docking is a less computationally expensive technique commonly employed for screening large libraries of small molecules against structured proteins with well-defined binding sites (26–29). It is presently unclear, however, if existing molecular docking approaches are suitable for characterizing IDP-ligand binding interactions or predicting the relative affinities of IDP ligands.

Traditional molecular docking methods aim to produce a single bound pose and can fail to account for protein conformational heterogeneity required for some protein-ligand binding events (30). To address this issue, flexible docking (31–34) and ensemble docking methods have been developed (30, 35–37). In flexible docking approaches, a subset of ligand and protein degrees of freedom, such as protein sidechain or ligand dihedral angles, are sampled during a docking calculation to search for alternative conformations that produce a better docking score. In ensemble docking protocols, docking is performed on conformational ensembles containing multiple protein conformations, which can be obtained from computer simulations or from experimental structures determined with different ligands bound.

Flexible docking and ensemble docking approaches have largely been applied to capture relatively subtle conformational changes in the binding sites of folded proteins, such as changes in sidechain rotamers and fluctuations in loop regions of otherwise well-defined 3D structures (38). It is currently unclear if existing ensemble docking approaches and docking scoring functions, which were developed to describe conformational fluctuations and ligand affinities in binding pockets of structured proteins, are well-suited to describe the dynamic and heterogenous binding mechanisms of IDP ligands.

Here, we propose two easily parallelized and computationally efficient ensemble docking protocols to predict the binding modes of small molecules to IDPs and asses their ability to predict the relative binding affinities of small molecules to α-synuclein, an extensively characterized IDP whose aggregation is associated with neuronal death in Parkinson’s disease (11, 39–40). We assess the ability of the proposed ensemble docking approaches to i) predict the relative binding affinities of α-synuclein ligands measured by solution NMR spectroscopy ii) reproduce the atomic-resolution details of ligand binding modes observed in long timescale all-atom MD simulations run with state-of-the-art force fields.

We test the performance of the ensemble docking protocols using AutoDock Vina (28, 34), a traditional force-field based molecular docking program, and DiffDock (29), a more recently developed deep learning approach based on a denoising diffusion generative model. We find that the proposed ensemble docking protocols correctly predict the relative affinities of the IDP ligands measured by NMR spectroscopy and reproduce the binding modes observed in long timescale MD simulations with remarkable accuracy. The ensemble docking protocols proposed here could provide a valuable, computationally efficient tool to study IDP ligand binding modes and discover novel IDP inhibitors.

## Results

We propose and validate two ensemble docking protocols for characterizing the dynamic and heterogenous binding modes of small molecules to IDPs. Experimental and computational studies have demonstrated that IDPs populate a heterogenous ensemble of rapidly interconverting conformations in solution and that small molecule IDP inhibitors discovered thus far do not stabilize a small subset of conformational states populated in apo IDP conformational ensembles (10–18, 22–25). Effective IDP ensemble docking protocols therefore require physically realistic conformational ensembles of IDPs that accurately reflect the populations of conformational states in solution as an initial input. Recent improvements in molecular mechanics force fields and water models have dramatically improved the accuracy of MD simulations of IDPs as assessed by their agreement with a large variety of experimental measurements (41–43). IDP ensembles obtained from long timescale or enhanced sampling MD simulations performed with modern force fields and validated or refined (9) with experimental measurements from solution NMR spectroscopy are therefore a natural choice for IDP ensemble docking methods.

We assess the ability of IDP ensemble docking protocols to accurately rank the experimental binding affinities of IDP ligands and reproduce atomic-level binding mechanisms observed in long timescale all-atom MD simulations of a previously studied C-terminal fragment of α-synuclein (11). The small molecule Fasudil was previously found to be neuroprotective in mouse models of Parkinson’s disease and shown to interact with monomeric α-synuclein by NMR chemical shift perturbations (CSPs) (39). Long timescale explicit solvent MD simulations, performed with the a99SB-*disp* protein force field and water model and the generalized amber force field (GAFF1) (41, 44), were subsequently used to study the binding mechanism of Fasudil to monomeric α-synuclein (11). A 1.5 millisecond unbiased MD simulation of full-length α-synuclein identified a 20-residue C-terminal fragment (residues 121–140) as having the highest the propensity to bind Fasudil, consistent with experimental NMR measurements (11). A 100μs MD simulation of this fragment (termed “α-syn-C-term”) was performed in its apo state and a 200μs MD simulations of α-syn-C-term were performed in the presence Fasudil to obtain a detailed statistical description of the heterogenous binding modes of Fasudil and to assess differences in the apo and holo (ligand-bound) conformational ensembles of α-syn-C-term. This study, and subsequent analyses (45), found the apo and holo ensembles of α-syn-C-term to be largely indistinguishable. MD simulations of α-syn-C-term were subsequently performed with 49 additional small molecules, and the binding affinities of five small molecules were characterized with NMR CSPs (11). The relative binding affinities of the ligands observed in MD simulations agreed with the experimental affinities measured by NMR.

Here, we perform ensemble docking calculations on α-syn-C-term using Fasudil and the highest affinity ligand (Ligand 47) and lowest affinity ligand (Ligand 23) previously characterized by NMR spectroscopy and MD simulations (Figure 1) (11). For each ligand, we use two ensemble docking approaches to calculate a heterogenous ensemble of docked poses, which we refer to as a *docked ensemble*. The first approach uses Autodock Vina (28), a traditional force-field based molecular docking program. The second approach uses DiffDock (29), a recently developed deep learning docking approach based on a denoising diffusion generative model. We compare the distributions of docking scores obtained from docked ensembles of each ligand and observe that both approaches correctly rank the relative binding affinities of the ligands determined by experimental NMR measurements.

**Figure 1.**
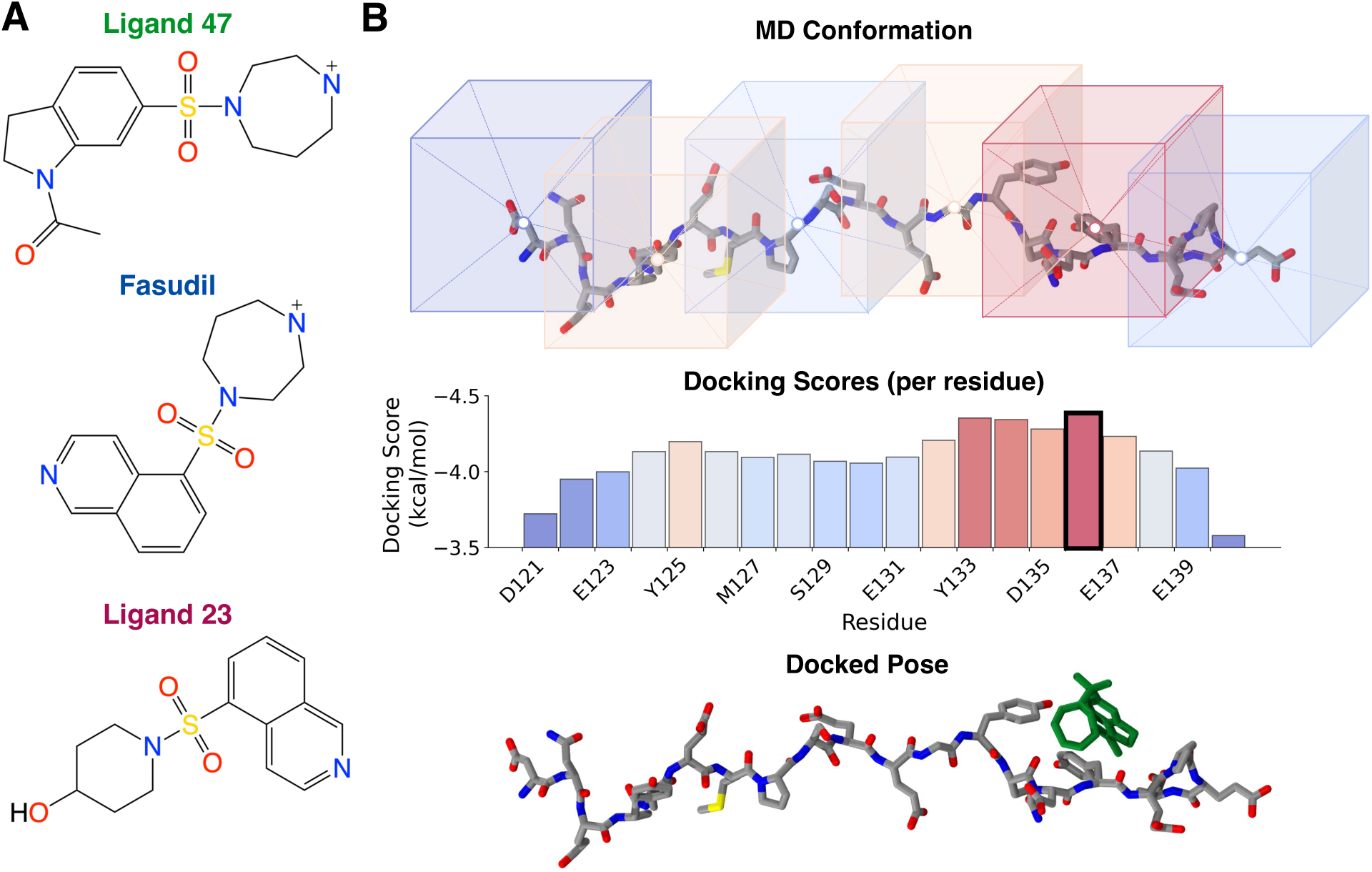
Ensemble docking for intrinsically disordered proteins. **(A)** A series of ligands that bind α-synuclein with experimental affinities previously determined by NMR spectroscopy (11). **(B)** A schematic illustration of the AutoDock Vina ensemble docking protocol for IDPs proposed here. For each conformation in an IDP ensemble, docking calculations are performed restricting the docking search space to a cubic volume surrounding the center-of-mass of each residue. One docked pose is returned for each residue, and the docked pose with the best docking score is selected as the docked pose for that conformation. This returns a *docked ensemble* containing one docked pose per conformation.

We provide detailed comparisons of the ligand binding modes present in docked ensembles obtained with AutoDock Vina and DiffDock and ligand-bound ensembles obtained from long timescale MD simulations. To enable a fine-grained comparison of docked ensembles and MD ensembles, we use a recently developed t-stochastic neighbor embedding (t-SNE) clustering method (45) to cluster α-syn-C-term conformations sampled in MD simulations into 20 conformational states. We compare the distribution of ligand binding modes and the populations of protein-ligand interactions in each α-syn-C-term conformational state across MD ensembles and docked ensembles and find remarkable agreement. We find that AutoDock Vina and DiffDock generate largely similar docked ensembles for α-syn-C-term but also identify some key differences.

### Ensemble docking protocols for intrinsically disordered proteins

The ensemble docking protocols proposed here require an ensemble of physically realistic conformations of an IDP as an input. We test the proposed ensemble docking protocols using ensembles of α-syn-C-term, a 20-residue fragment consisting of residues 121-140 of the IDP α-synuclein, obtained from previously reported explicit-solvent all-atom MD simulations performed in its apo state and in the presence of ligands (11). MD simulations of α-syn-C-term performed in its apo state and in the presence of the ligands studied here (Fasudil, Ligand 47 and Ligand 23) were found to be in excellent agreement with NMR backbone chemical shifts, scalar couplings, and residual dipolar couplings (RDCs), demonstrating that the simulated α-syn-C-term conformational ensembles are accurate descriptions of the solution ensemble of this IDP (11).

We perform docking calculations on an α-syn-C-term ensemble obtained from an apo MD simulation, which we refer to as *apo docking,* to assess predictive power of ensemble docking to prospectively rank potential binders without using prior information from ligand binding MD simulations. We also perform *holo docking* calculations, which are sometimes referred to as *redocking* calculations, to directly compare ligand binding poses and the statistical properties of ligand-bound ensembles obtained from MD and ensemble docking. In holo docking calculations we first obtain ligand-bound α-syn-C-term ensembles from MD simulations of α-syn-C-term with each ligand (Fasudil, Ligand 47, or Ligand 23) and remove the ligand coordinates before performing docking. We define a ligand-bound MD ensemble as all frames in an MD simulation where at least one heavy (non-hydrogen) atom of α-syn-C-term is within 6Å of at least one heavy ligand atom.

We perform ensemble docking calculations on conformational ensembles of α-syn-C-term using two approaches. In both approaches, we perform docking calculations on each conformation, or *frame*, of an input ensemble and return one docked pose for each frame to produce a *docked ensemble*. In the first ensemble docking approach, we perform multiple docking calculations on conformation frame using AutoDock Vina, a popular open-source docking method that employs a physics-based scoring algorithm (28). For each frame in the input ensemble we perform 20 docking calculations, restricting the docking search space to a cubic volume surrounding the center-of-mass of one residue, and return one candidate docking pose for each residue. We evaluate the docking score of each predicted pose of each frame using the AutoDock Vina scoring function and select the best scoring pose of the 20 predicted poses as the final predicted docked pose that frame. This approach is schematically illustrated in Figure 1B. We subsequently refer to this approach as “AutoDock Vina ensemble docking”. Further details are provided in “AutoDock Vina ensemble docking” in the Methods section.

In the second ensemble docking approach, we perform one docking calculation on each conformation in the input ensemble with no restrictions on the docking search space and allow the docking algorithm to identify the optimal binding pose of each frame. For these calculations we use the recently developed DiffDock method (29), which uses a denoising diffusion generative model to efficiently search for an optimal ligand binding pose for an input structure and reports a *confidence score* for each docked pose. We subsequently refer to this approach as “DiffDock ensemble docking”. Further details are provided in “DiffDock ensemble docking” in the Methods section. We compare the results of both docking approaches in apo docking and holo docking calculations.

To obtain robust statistics of docking scores and the properties of docked ensembles obtained with each ensemble docking approach, we perform apo docking calculations on an apo α-syn-C-term MD ensemble containing 20,000 conformations. To facilitate detailed comparisons of the ligand binding poses predicted by ensemble docking and poses observed in unbiased MD simulations, we use a recently developed t-SNE clustering algorithm (45) to cluster each α-syn-C-term MD ensemble into 20 conformational states. We compare the ligand binding modes observed in MD ensembles and docked ensembles in each of the 20 conformational states. To obtain statistically equivalent comparisons of docked poses and MD poses in each conformational state, we perform docking calculations on 1000 conformations, randomly selected without replacement, from each t-SNE cluster of α-syn-C-term. Holo docking calculations of Fasudil and Ligand 47 were performed on ensembles of 20,000 α-syn-C-term conformations (20 t-SNE clusters containing 1000 conformations each). Holo docking calculations of Ligand 23 were performed on an ensemble of 18,861 conformations, as not all t-SNE clusters obtained from the MD simulation of Ligand 23 binding contained 1000 ligand-bound conformations. Since the t-SNE clusters identified from MD simulations have unequal populations, we weight the average values of ensemble properties (such as normalized docking scores or intermolecular interaction populations) by the population of each t-SNE cluster from the initial unbiased MD ensemble used as input for docking

### Ensemble docking accurately predicts the relative affinities of small molecules to α-synuclein

To predict the relative affinities of α-synuclein ligands with ensemble docking we perform apo and holo docking calculations with AutoDock Vina and DiffDock for Fasudil, Ligand 47 and Ligand 23 and compute the docking scores of every frame in each docked ensemble. We calculate the average docking score of each docked ensemble (Figure 2, SI Table 1). To enable comparisons of relative affinity predictions from AutoDock Vina and DiffDock we normalize the docking scores of all ligands obtained with each method to a scale of 0 to 1 using the maximum and minimum docking score values observed in calculations of all three ligands with each docking approach (“Comparing docking scores” in Methods, Eq. 1). For each ensemble docking approach tested (AutoDock Vina holo docking, AutoDock Vina apo docking, DiffDock holo docking, DiffDock apo docking) we use one normalization scale for the docking scores of all three ligands. For each ensemble docking approach, a normalized docking score of 1 is defined as the most favorable docking score observed in docking calculations of all three ligands, and a normalized docking score of 0 is defined as the least favorable docking score observed in all docking calculations (Eq. 1). Uncertainties of normalized docking scores were calculated from bootstrapping using 10,000 samples for each docked ensemble (SI Table 1).

**Figure 2.**
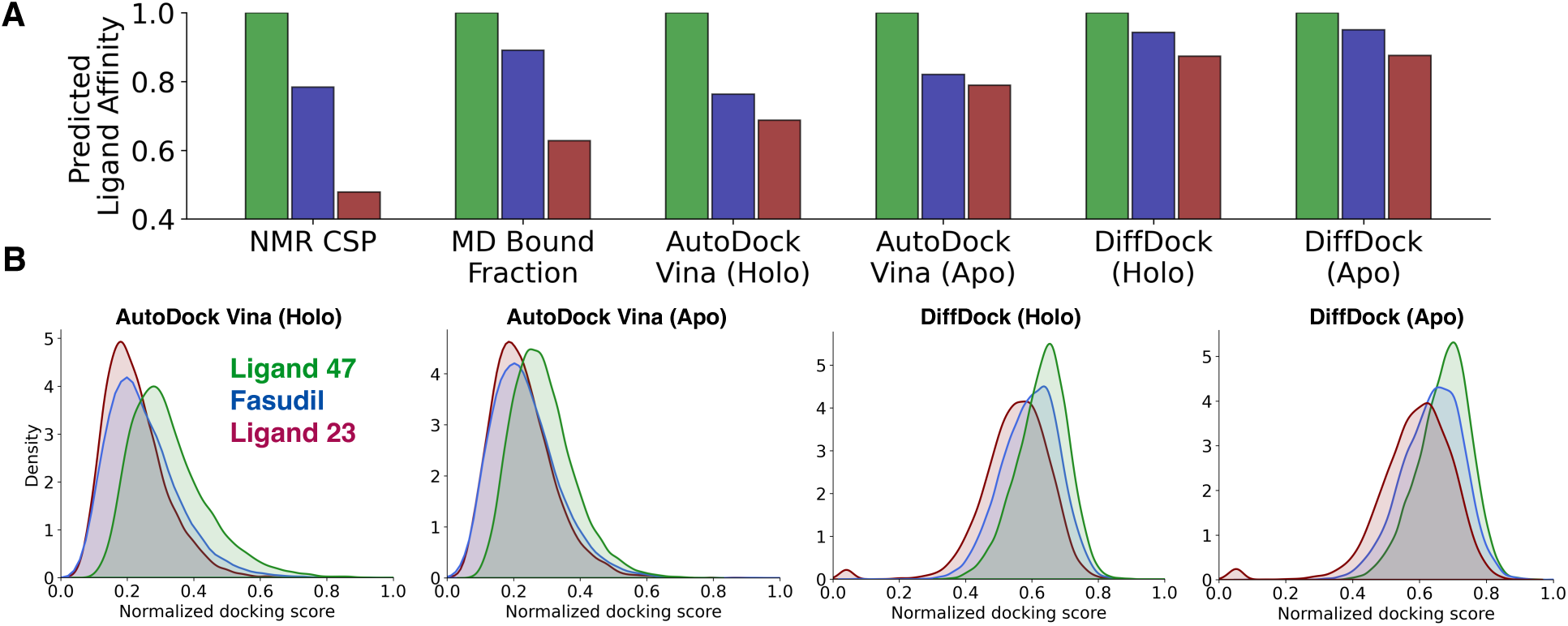
IDP ligand affinity predictions from ensemble docking are consistent with experimental affinities from NMR spectroscopy and simulated affinities from MD simulations. **(A)** Comparison of the relative ligand affinities predicted by the proposed ensemble docking protocols with experimental ligand affinities determined from NMR chemical shift perturbations (CSPs) and affinity predictions from long timescale MD simulations (11). Relative experimental affinities from NMR and affinity predictions from MD and each docking approach are scaled such that tightest binding ligand has a relative affinity value of 1.0. **(B)** Distributions of normalized docking scores from holo and apo Autodock Vina and DiffDock ensemble docking calculations. Docking scores obtained from each docking method have been normalized with min-max normalization using the highest and lowest docking score observed in calculations of all three ligands.

We compare the average normalized docking scores obtained for each ligand with each ensemble docking approach with the relative binding affinities determined from NMR spectroscopy and the relative binding affinities observed in long timescale unbiased MD simulations of α-syn-C-term in the presence of each ligand in Figure 2A. MD binding affinities are defined as the fraction of frames in the MD simulation where at least one heavy atom of α-syn-C-term is within 6Å of at least one heavy atom of the ligand. NMR binding affinities were determined based on the relative magnitudes of NMR CSPs for the backbone amide groups of α-synuclein residues Y125, Y133 and Y136 (11). To enable comparisons with docking predictions, we normalize the relative ligand binding affinities measured by NMR and calculated from unbiased MD simulations to the binding affinity of Ligand 47, the highest affinity binder. All ensemble docking approaches predict the correct rank order of ligand affinities measured by NMR and predicted by long-time scale MD simulation. Holo docking with AutoDock Vina predicts the greatest difference in average normalized docking score between Ligand 47 and Fasudil, whereas apo docking with DiffDock produces more similar average normalized docking scores for Ligand 47 and Fasudil (Figure 2, SI Table 1).

To gain a more detailed understanding of the distribution of docking scores within each docked ensemble, we compute the average normalized docking score of each cluster identified by t-SNE clustering (SI Figure 1). We characterize each t-SNE cluster by the angle formed by the Ca atoms on residues 121, 131 and 140 of α-syn-C-term, which we refer to as the “bend angle” of the protein fragment (45). The relative ligand affinity predictions are largely consistent across clusters and that the ensemble averaged docking scores of each ligand are not dictated by outliers in a small number of clusters. We observe strong correlations between the average α-syn-C-term bend angle and average normalized docking score of t-SNE clusters in each ligand (SI Figure 1, SI Table 2). Higher docking scores are observed for smaller α-syn-C-term bend angles, which are associated with more compact conformations that enable the simultaneous formation of contacts with multiple regions of α-syn-C-term, in all ensemble docking approaches. There is a smaller correlation between bend angle and docking scores in DiffDock calculations than AutoDock calculations, and the correlation is particularly weak for DiffDock calculations of the Ligand 23, the lowest affinity ligand (SI Figure 1). We observe that the correlations between docking score and bend angle are substantially larger than the correlations between the MD bound fraction and bend angle (SI Figure 2, SI Table 2). This suggests that the AutoDock Vina docking score and DiffDock confidence score favor binding to compact conformations over extended conformations more so than explicit solvent MD ligand binding simulations

### Ensemble docking accurately reproduces IDP ligand binding modes observed in experimentally validated long timescale MD simulations

To assess the similarity of IDP ligand binding modes obtained from ensemble docking and from MD simulations, we perform holo docking calculations of Fasudil, Ligand 47 and Ligand 23 to α-syn-C-term with the proposed AutoDock Vina and DiffDock ensemble docking approaches. For each ensemble docking calculation, we compare the similarity of the ensemble of docked poses and ligand-bound poses from MD in each of the 20 α-syn-C-term conformational states identified by t-SNE clustering (SI Tables 3-4, SI Figures 3-6). For each cluster identified by t-SNE, we compute the populations of protein-ligand contacts and the populations of specific protein-ligand intermolecular interactions (hydrogen bonds, charge contacts, aromatic stacking interactions, and hydrophobic contacts) between docked ligands and each residue of α-syn-C-term as described in the “Analysis of IDP ligand binding modes” section in methods (Figure 3, SI Figures 3-4). To assess the cooperativity of intermolecular interactions involved in ligand binding obtained with each approach, we calculate the probability that a ligand simultaneously forms contacts with a pair of residues in α-syn-C-term, which we refer to as a *dual-residue contact probability* (Figure 3, SI Figures 5-6).

**Figure 3.**
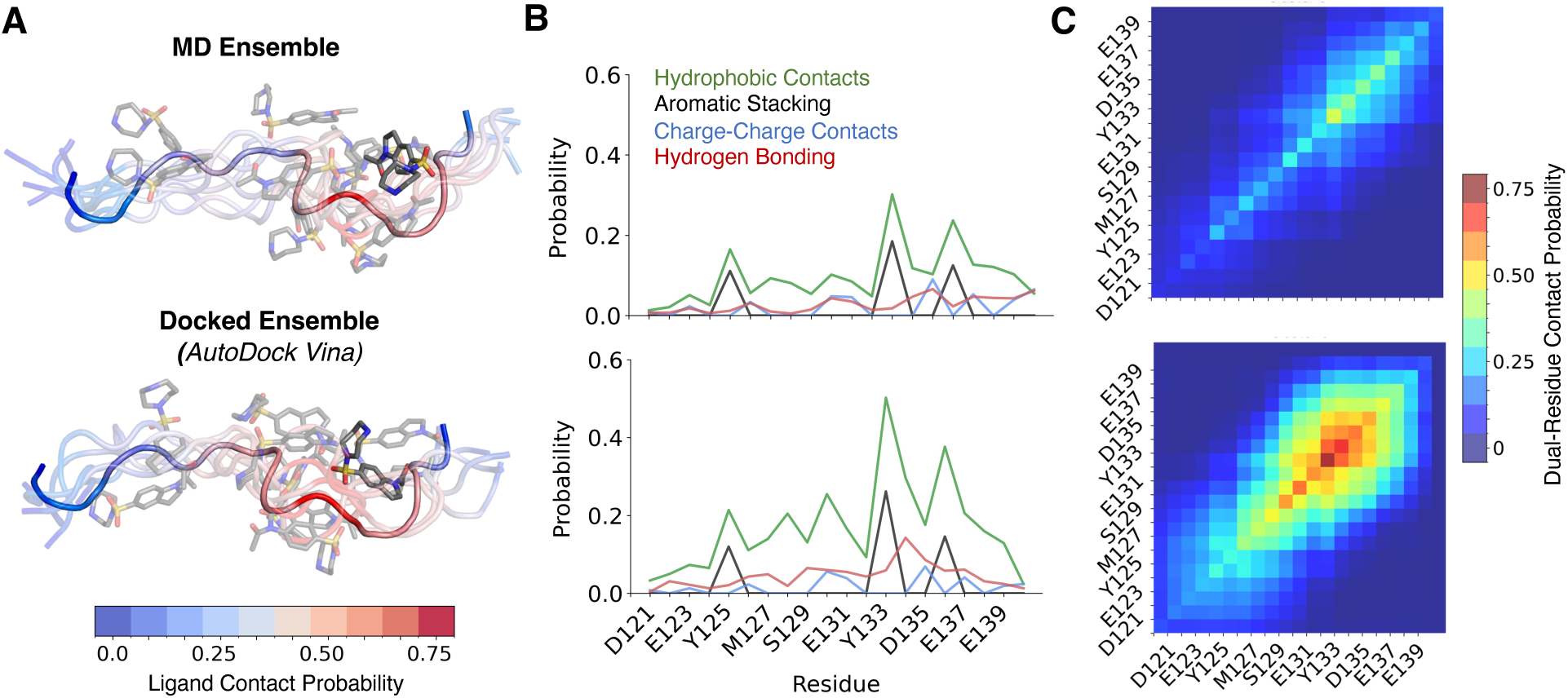
Comparison of ligand binding poses obtained from long timescale MD simulations and IDP ensemble docking calculations. Comparisons of Ligand 47 binding modes obtained from a long timescale MD simulation (top) and AutoDock Vina holo ensemble docking (bottom) for a representative conformational substate of α-syn-C-term identified by t-SNE clustering. The MD ligand-bound ensemble contains all frames from an unbiased MD simulation where at least one heavy atom of α-syn-C-term is within 6Å of at least one heavy atom of the ligand. In holo docking calculations, the ligand is removed from all conformations in the ligand-bound MD ensemble, and a docked pose is predicted for each conformation. **(A)** Overlay of representative snapshots of ligand-bound conformations from MD and from ensemble docking in the selected t-SNE cluster. α-syn-C-term residues are colored by a gradient corresponding to the contact probability of Ligand 47 with each residue in the selected cluster. **(B)** Populations of intermolecular interactions between Ligand 47 and each residue of α-syn-C-term in ligand-bound ensembles in the selected t-SNE cluster. **(C)** The probability that Ligand 47 simultaneously forms contacts with each pair of residues of α-syn-C-term residues in the selected t-SNE cluster.

A comparison of the ensemble of Ligand 47 binding poses obtained from AutoDock Vina holo docking and from a 200μs MD simulation of Ligand 47 binding is shown for a representative α-syn-C-term cluster (Cluster 0, which has an average α-syn-C-term bend angle of 148°) in Figure 3. A visual comparison of subsets of Ligand 47 bound poses from the docked ensemble and the MD ensemble are shown in Figure 3A. The per-residue populations of protein-ligand contacts observed in the docked ensemble and the MD ensemble have a Pearson correlation coefficient (*r*) of *r*=0.91 in this cluster. In Figure 3B we compare the per-residue populations of specific intermolecular interactions in the docked ensemble and MD ensemble in this cluster. The populations of hydrophobic contacts, aromatic stacking interactions, charge contacts and hydrogen bonds observed in the docked ensemble and MD ensemble have correlation coefficients of *r*=0.93, *r*=1.00, *r*=0.81 and *r*=0.42, respectively, in this cluster. We compare the dual-residue contact probabilities obtained from ensemble docking and from MD in Figure 3C. We observe that the dual-residue contact populations of the docked ensemble and MD ensemble are highly correlated (*r* =0.94), but the dual-residue contact probabilities in docked ensembles are systematically larger (*RMSE* = 0.14).

We observe close agreement between the populations of Ligand 47 interactions and dual-residue contact probabilities in the AutoDock Vina holo docked ensembles and MD ensembles across all 20 t-SNE clusters (SI Figures 3-6, SI Tables 3-4). The per-residue populations of hydrophobic contacts, aromatic stacking interactions, and charge interactions are similar between the MD and AutoDock Vina holo docked ensembles across all t-SNE clusters, with average correlation coefficients of r = 0.90, r = 0.88, and r = 0.76, respectively. The correlation for hydrogen bond populations is weaker, with an average correlation coefficient of r = 0.39. We observe a strong correlation of the populations of dual-residue contacts averaged across all t-SNE clusters (r=0.90) and draw attention to the striking similarity in the variations of patterns of dual-residue contacts across t-SNE clusters (SI Figures 5-6). The dual-residue contact analysis demonstrates that AutoDock Vina does not predict the same distribution of binding modes for all clusters, and instead faithfully reproduces differences in the distributions of binding modes present in MD.

The ensemble averages of the populations of Ligand 47 intermolecular interactions observed in MD and AutoDock Vina holo docking are in close agreement (Figure 4, SI Table 4). The ensemble averaged per-residue populations of hydrophobic contacts, aromatic stacking interactions, charge contacts, hydrogen bonds have correlation coefficients of *r*=0.91, *r*=0.94, *r*=0.97 and *r*=0.51, respectively, and the ensemble averaged populations of dual-residue contacts have a correlation coefficient of r=0.94. We observe similarly close agreement between Autodock Vina holo docked ensembles and MD simulations for Fasudil and Ligand 23 (SI Tables 3-4).

**Figure 4.**
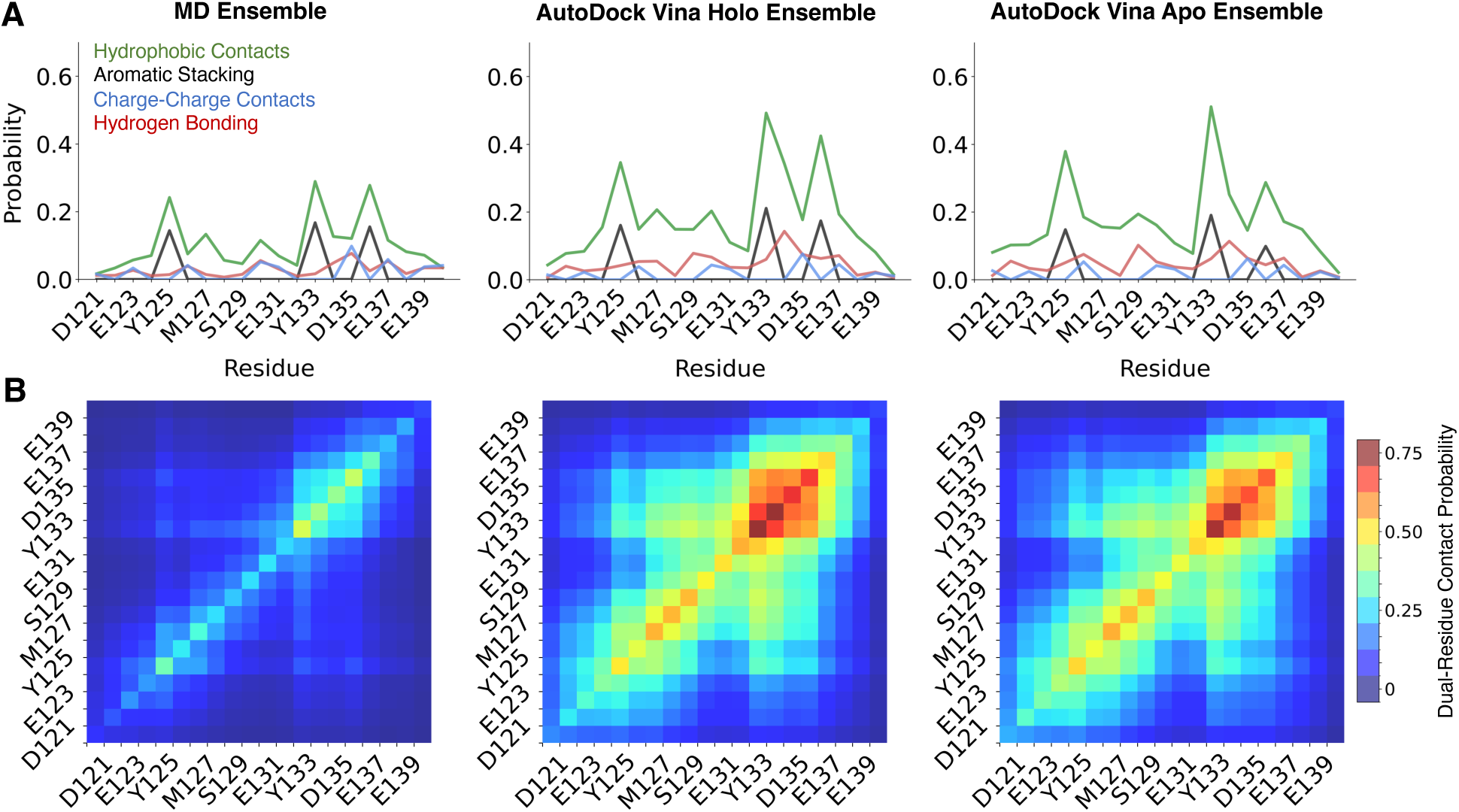
Comparison of ensemble averaged intermolecular protein-ligand interactions and dual-residue contact probabilities between Ligand 47 and α-syn-C-term obtained from a 200μs MD simulation with Ligand 47, AutoDock Vina holo docking calculations, and AutoDock Vina apo docking calculations. **(A)** Average populations of intermolecular interactions between Ligand 47 and each residue of α-syn-C-term in ligand-bound ensembles, weighted by the populations of each cluster identified by t-SNE for the original MD simulation. **(C)** The average probabilities that Ligand 47 simultaneously forms contacts with each pair of residues in α-syn-C-term, weighted by the populations of each cluster identified by t-SNE for the original MD simulation.

Having established that holo docking (or redocking) calculations performed with Autodock Vina are in excellent agreement with long timescale MD, we proceed to compare the docked ensembles obtained from the more realistic scenario of apo docking, where different ligands are docked onto the same apo α-syn-C-term MD ensemble. We compare the populations of intermolecular interactions and dual-residue contacts between Ligand 47 and α-syn-C-term in ligand-bound ensembles obtained from MD, AutoDock Vina holo docking, and AutoDock vina apo docking in Figure 4. We observed that the Ligand 47 docked ensembles obtained from AutoDock Vina apo docking and holo docking results are highly similar (SI Figures 7-8, SI Tables 3-4).

We compare the Fasudil and Ligand 23 docked ensembles obtained from AutoDock Vina apo docking and holo docking in SI Figures 9-10. Docked ensembles of Fasudil obtained from AutoDock Vina are in close agreement with MD. The agreement between AutoDock Vina docked ensembles and MD ensembles is substantially worse for Ligand 23 (SI Tables 3-4, SI Figure 10), the lowest affinity α-synuclein ligand studied here. Ligand 23 lacks the positively charged amine group shared by Ligand 47 and Fasudil, suggesting that the charge contacts between the amine groups of these ligands and the negatively charged aspartate glutamate acid sidechains of α-syn-C-term are important for recovering ligand binding poses similar to those observed in MD.

The apo and holo AutoDock Vina docked ensembles of Fasudil and Ligand 23 are very similar (SI Figures 9-10). We observe that while the populations of aromatic stacking and charge interactions observed in apo docked ensembles and holo ensembles have similar deviations from the populations observed in MD simulations, the rank order of the per-residue populations of these interactions aligns more closely with MD in holo docked ensembles, leading to significantly higher correlation coefficients (SI Tables 3-4). This suggests that while there is almost no detectable difference in the distribution of backbone conformations in apo and holo α-syn-C-term ensembles, the binding of each ligand influences the relative orientations of α-syn-C-term sidechain orientations in MD holo ensembles strongly enough to affect the results of docking calculations. The relative positions of sidechain pharmacophores in MD holo ensembles and apo ensembles produce globally similar docked ensembles with subtle differences in the distributions of binding poses and populations of intermolecular interactions (SI Figure 4, SI Figures 6-10).

We compare the results of DiffDock ensemble docking calculations to MD ensembles and AutoDock Vina docked ensembles (SI Tables 3-4, SI Figures 11-17). There is reasonably good agreement between ligand-bound ensembles obtained from DiffDock and MD, but we observe some notable differences between DiffDock and AutoDock Vina docked ensembles. The populations of intermolecular interactions in DiffDock docked ensembles have substantially lower correlation coefficients with ligand-bound ensembles obtained from MD (SI Tables 3-4). DiffDock docked ensembles have substantially lower populations of all intermolecular interactions other aromatic stacking, and the populations of charge contacts and hydrogen bonds are close to zero for most residues (SI Figures 11-12). This suggests that these interactions do not have a strong effect on the orientations of IDP-ligand binding poses predicted by DiffDock.

Nearly all intermolecular interactions populated in DiffDock ensembles are made with the three aromatic residues of α-syn-C-term (Y125, Y133 and Y136) and their immediate neighbors. This suggests that aromatic interactions are the dominant feature guiding DiffDock binding pose predictions in this system. The distribution of binding poses obtained from DiffDock has substantially less variation among t-SNE clusters relative to AutoDock (SI Figures 14-15), where a diversity of cluster-dependent patterns of dual-reside contacts are observed (SI Figure 6, SI Figure 8). This suggests that the results of AutoDock Vina calculations may be more sensitive to differences in IDP conformations than DiffDock. AutoDock Vina ensemble docking systematically overestimates the populations of intermolecular interactions relative to MD ensembles, while DiffDock ensemble docking systematically underestimates the populations of all intermolecular interactions other than aromatic stacking. AutoDock Vina overestimates the populations of dual-residue contacts relative to MD, but more faithfully capture the pattern of dual-residue contacts observed in MD simulations that DiffDock (SI Figures 5-6, SI Figure 8, SI Figures 14-15).

### Quantifying the similarity of ligand binding poses obtained from MD simulations and ensemble docking

We compare the similarity of the individual binding poses obtained from ensemble docking and MD by computing the root mean square deviation (RMSD) of ligand poses predicted from docking and ligand poses observed in MD simulations (Figure 5). We consider two RMSD metrics. To analyze ensembles from holo docking, where there is a reference ligand-bound MD pose for each α-syn-C-term conformation used for docking, we directly compute the RMSD between the ligand heavy atom coordinates of the docked pose and the original MD pose after aligning on only protein coordinates. We refer to this value as a *frame-matched RMSD.* We display the distribution of frame-matched RMSD values for Ligand 47 holo docking calculations in Figure 5A.

**Figure 5.**
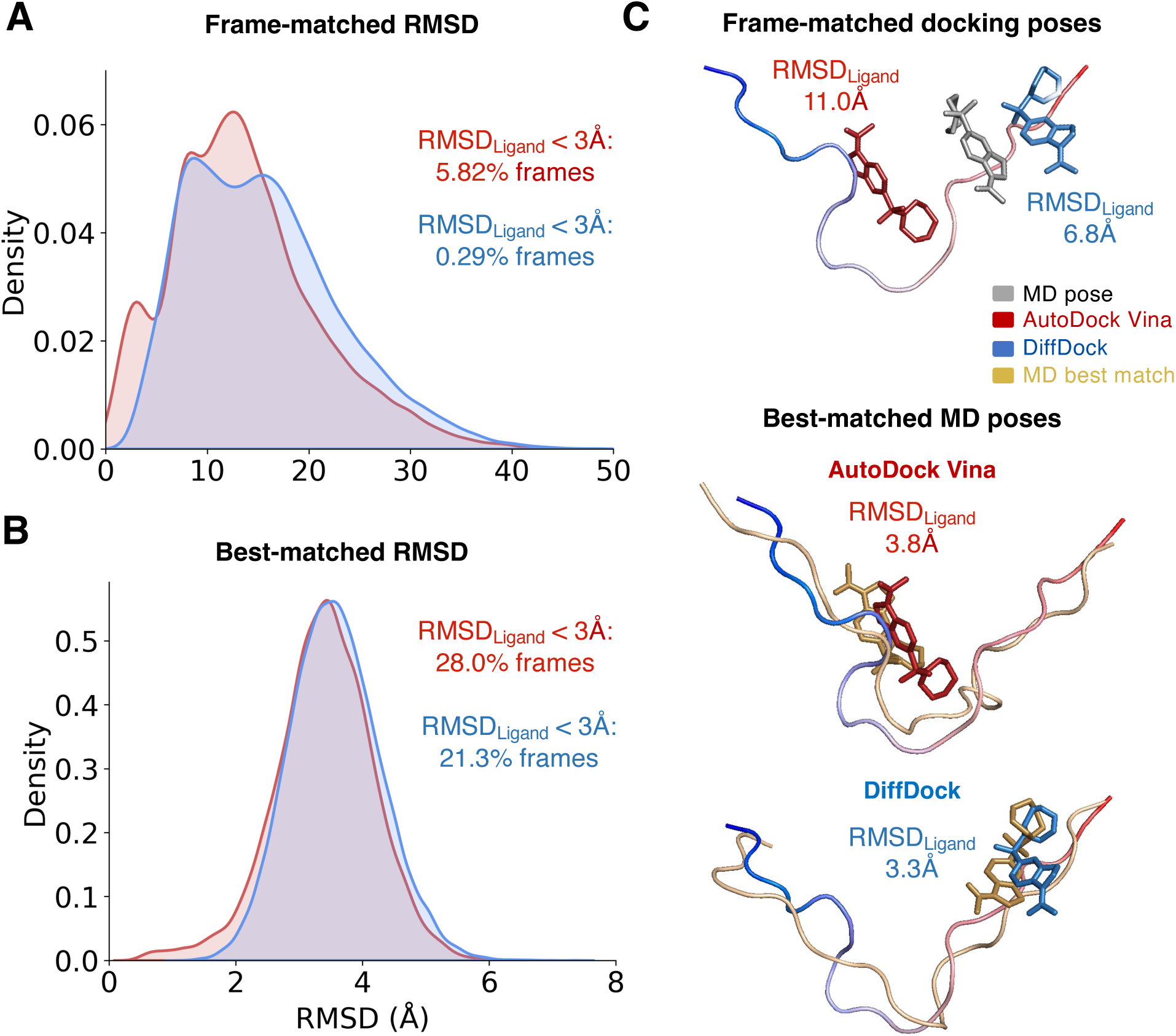
Comparison of the ligand RMSD between docked poses and MD bound poses pf Ligand 47. Distribution of frame-matched RMSD values **(A)** and best-matched RMSD values **(B)** obtained from AutoDock Vina holo docking calculations (red) and DiffDock holo docking calculations (blue) of Ligand 47. The fraction of frames with ligand RMSD values less than 3Å are displayed for each docking method. **(C)** A representative frame showing the α-syn-C-term protein coordinates used for docking in a blue-to-red gradient. The docked ligand poses predicted by AutoDock Vina ensemble docking (red) and by DiffDock docking (blue) are compared to the ligand pose in the original MD bound pose (gray), and the ligand RMSD values between the docked pose and the MD pose are displayed (top). The best-matched MD poses are shown for the AutoDock Vina docked pose (middle) and DiffDock docked pose (top). The α-syn-C-term coordinates and ligand coordinates of the best-matched MD poses are colored tan. The ligand RMSD values between the best-matched MD pose and each docked pose are displayed.

We observe that AutoDock Vina and DiffDock rarely predict bound poses similar to the original MD pose. Only 5.8% of Autodock Vina pose predictions and 0.3% of DiffDock pose predictions have a ligand RMSD less than 3.0Å. Only 10.7% of Autodock Vina predictions and 3.6% of DiffDock pose predictions have a ligand RMSD less than 5.0Å. The fraction of frames with ligand RMSDs beneath these thresholds is even lower in docked ensembles of Fasudil and Ligand 23 (Table 1). Most holo docking poses predict identify entirely different binding sites than the corresponding MD frame, with ligand RMSDs greater than 10 Å (Figure 5C). This result is however, relatively unsurprising as the ligand binding mechanisms observed in MD simulations of α-syn-C-term were found to be highly diffusive, with no identifiable one-to-one mappings between protein conformations and ligand binding poses (11, 45). Many distinct ligand poses were found to have similar populations in each conformational substate of α-syn-C-term (45).

**Table 1.**
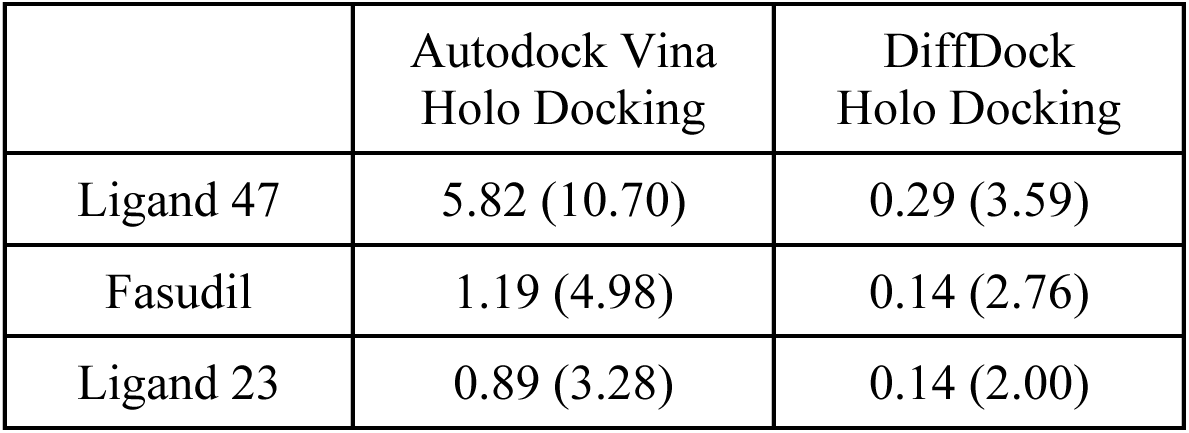
Similarity of frame-matched ligand RMSD of ligand poses predicted by docking and reference poses observed in MD simulations. We report the percentage of docked frames obtained by holo ensemble docking where the frame-matched RMSD of ligand heavy-atom coordinates is less than 3Å from the original MD bound-pose and where the frame-matched RMSD is less than 5Å from the original MD bound-pose. The percentages of frames with a frame-matched ligand RMSD less than 5Å are reported in parentheses.

Accordingly, we compute a second RMSD metric, the *best-matched RMSD*. For each docked pose, we align the protein coordinates of all bound poses observed in the ligand-bound MD ensemble to the protein coordinates of the docked pose and compute the ligand RMSD (Figure 5C). We identify the MD frame with the smallest ligand RMSD as the best-matched MD frame and report the value of the ligand RMSD as the best-matched RMSD for this pose. This allows us to quantify the deviation of the ligand position of each docked pose to the most similar ligand-bound pose observed in long timescale MD simulations. We display the distribution of best-matched RMSD values for Ligand 47 holo docking calculations in Figure 5B and report the fraction of docked poses with best-matched ligand RMSDs less than 3.0Å and 5.0Å in Table 2.

**Table 2.**
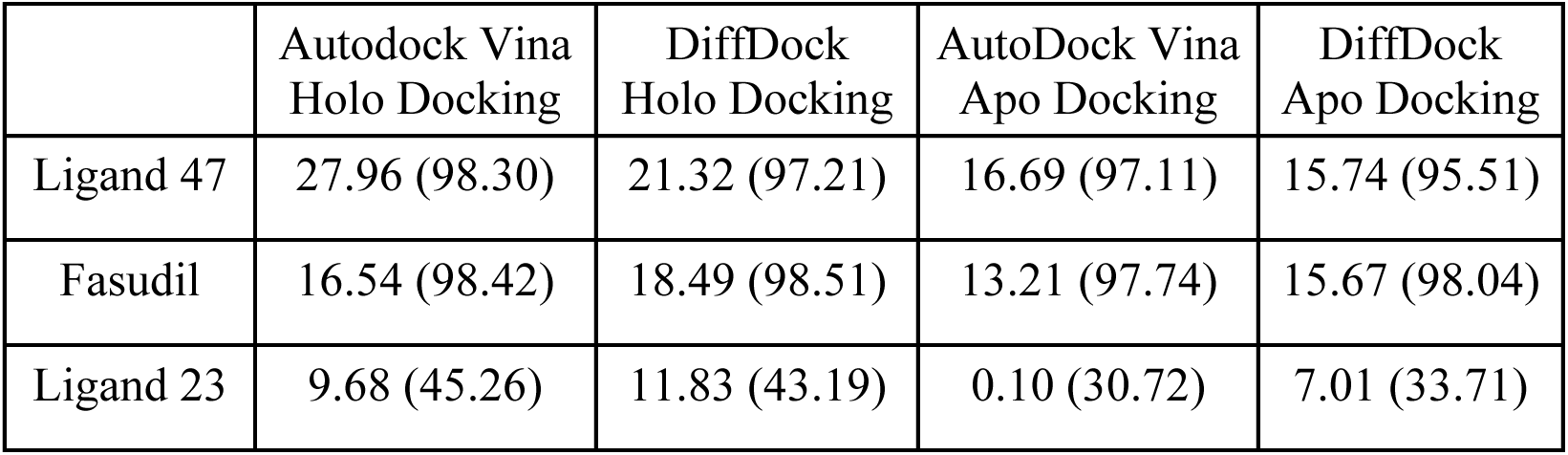
Similarity of best-matched ligand RMSD of ligand poses predicted by docking and reference poses observed in MD simulations. We report the percentage of docked frames obtained by ensemble docking where the best-matched RMSD of ligand heavy-atom coordinates is less than 3Å from the original MD bound-pose and where the best-matched RMSD is less than 5Å from the original MD bound-pose. The percentages of frames with a best-matched ligand RMSD less than 5Å are reported in parentheses. Reference ligand poses for apo docking ensembles were obtained from the long-time scale MD simulation of α-syn-C-term with the docked ligand.

|  | Autodock Vina<br>Holo Docking | DiffDock<br>Holo Docking | AutoDock Vina<br>Apo Docking | DiffDock<br>Apo Docking |
| --- | --- | --- | --- | --- |
| Ligand 47 | 27.96 (98.30) | 21.32 (97.21) | 16.69 (97.11) | 15.74 (95.51) |
| Fasudil | 16.54 (98.42) | 18.49 (98.51) | 13.21 (97.74) | 15.67 (98.04) |
| Ligand 23 | 9.68 (45.26) | 11.83 (43.19) | 0.10 (30.72) | 7.01 (33.71) |

A substantially larger fraction of frames in docked ensembles have best-matched RMSDs less than 3.0Å and 5.0Å compared to frame-matched RMSDs. 28% of Ligand 47 poses obtained from AutoDock Vina holo docking have a best-matched ligand RMSD of less than 3.0Å. This value drops to 21% in the DiffDock holo docked ensemble. We compare the distributions of best-matched ligand RMSDs observed in docked ensembles of Fasudil and Ligand 23 in SI Figures 18-19. In all docked ensembles of Fasudil and Ligand 47, over 95.0% of docked poses have a best-matched RMSD of less than 5.0Å. These values are substantially lower for Ligand 23, demonstrating that ensemble produces a large fraction of conformations not sampled in MD for this lower affinity ligand (Table 2, SI Figures 18-19).

Lastly, we consider the scenario where the protein coordinates of a ligand-bound MD ensemble of one ligand are used as input for ensemble-docking calculations of a different ligand. This scenario, which we refer to as *cross docking*, could potentially be used to predict the affinity of ligands with a similar scaffold after performing computationally expensive MD simulations with an initial ligand. We compare the distribution of the best-matched RMSD values obtained from cross docking Ligand 47 on the protein coordinates of a holo MD ensemble of α-syn-C-term bound to Fasudil and from cross docking Fasudil on the protein coordinates of holo MD ensemble of α-syn-C-term bound to Ligand 47 in SI Figure 20 and SI Table 5. We compare the ensemble averaged docking scores obtained from cross-docking in SI Table 6. We observe that in both cross-docking scenarios, the ensemble averaged docking scores identify the ligand that α-syn-C-term was originally simulated with as the highest affinity binder, erroneously predicting Fasudil to have a higher affinity than Ligand 47 when the protein coordinates from a Fasudil-bound MD ensemble are used as starting structures for docking Ligand 47. This suggests that for ligands with small differences in docking scores, *cross docking* with the protocols tested here may not provide reliable affinity predictions.

## Discussion

The field of IDP drug discovery has gained substantial momentum in recent years. Several small molecules have been discovered that directly bind and inhibit the interactions of IDPs without stabilizing the formation of rigid structural elements. Several ligands that bind IDPs through heterogenous and dynamic binding mechanisms have been shown to have clear in vitro affinity, in vivo activity and therapeutic effects in animal models (10–18). Multiple IDP ligands with disordered binding mechanisms have now entered human trials (19–21). There has also been growing interest in pursuing the discovery of IDP ligands that modulate the properties of biomolecular condensates as a novel route for the discovery of therapeutic compounds (13, 46–48).

Families of small molecules that span a range of *in vitro* affinities and *in vivo* potencies have been discovered for several IDP targets of pharmaceutical interest including α-synuclein (11, 39), the androgen receptor (13, 16–17, 19, 24, 49–51), p53 (14, 52), hIAPP (15), Abeta42 (12, 23, 53), p27kip1 (10, 54), TDP-43 (55–56) and c-Myc (18, 22, 57–59). Studies combining computational methods, biophysical experiments, cellular assays, and animal models have elucidated structure-affinity-activity relationships in several families of IDP ligands, in some cases facilitating the design of more potent IDP ligands with heterogenous binding mechanisms (11, 13). Computational investigations of androgen receptor (13, 24, 49) and α-synuclein ligands (11, 45) employed all-atom MD simulations ranging from 60-1500μs to elucidate atomic resolution binding mechanisms that successfully rationalize the affinity and potency of ligands with similar scaffolds. This demonstrates that atomic resolution models of heterogenous IDP ligand binding mechanisms have the potential to provide valuable insights in IDP drug discovery campaigns. These simulations however, require months of simulation time on special purpose supercomputers or parallel multi-GPU compute architectures making them impractical for screening large libraries of ligands.

Molecular docking presents an intriguing possibility for studying IDP ligand binding mechanisms and predicting the affinity of IDP ligands with substantially higher throughput than all-atom MD simulations. It has, however, been unclear if the energy functions or AI models used to predict small molecule binding sites are suitable for studying IDP ligand binding or predicting physically realistic atomic resolution models of heterogenous binding events. Initial studies have explored the use of docking to identify potential IDP ligands (52, 59). Thus far, these studies have used a combination of clustering and ligand-cavity prediction tools to identify a small number of structures and binding sites to use for high-throughput screening in an analogous fashion to docking campaigns for folded proteins with structured binding sites. While these studies have succeeded in identifying molecules with *in vitro* affinity or *in vivo* activity, they did not attempt to enumerate ensembles of binding modes with a realistic conformational ensemble that reflects the full conformational diversity of IDPs in solution or predict the relative affinities of ligands.

In one previous study, an ensemble docking approach was proposed to identify potential binding sites of the promiscuous polyphenol ligand epigallocatechin gallate (EGCG) in the disordered N-terminal domain of p53 (p53-NTD) (14). The authors used AutoDock Vina to dock EGCG to a sparse MD ensembles containing 100 conformations obtained from a relatively short 500ns MD simulation initiated from an extended linear structure. The authors observed that that docked poses of EGCG had a higher probability of being located near aromatic residues compared to other residues, in agreement with experimental p53-NTD NMR chemical shift perturbations (CSPs) measured in EGCG titrations. However, they did not assess the accuracy or physical plausibility of the p53-NTD MD ensemble or the docked EGCG poses or develop a metric to predict the relative affinities of different ligands.

Here, we demonstrate the first application (to our knowledge) of molecular docking to successfully predict the relative affinities of small molecule ligands to a realistic, experimentally validated conformational ensemble of an IDP. We demonstrate that the ensemble docking approaches proposed here produce docked ensembles that are highly similar to ligand-bound ensembles obtained from long-timescale, well-converged MD simulations performed with a state-of-the-art force field and water model. Prior to this study, it has been unclear if the scoring functions used in molecular docking programs, which have largely been trained to predict the affinity of small molecules to rigid hydrophobic binding sites, would be applicable for studying the heterogenous binding mechanisms of small molecules to IDPs. Here, we have performed ensemble docking on what may be a difficult edge case for molecular docking scoring functions: a highly charged disordered IDP fragment that predominantly samples extended solvent-exposed conformations, and samples very few conformations that resemble the binding sites of folded proteins. We observe that even in this challenging case, ensembles of ligand-bound poses obtained from ensemble docking (particularly from ensemble docking performed with AutoDock Vina) are highly similar to MD ensembles. These results demonstrate that ensemble molecular docking strategies for IDPs have substantial potential and warrant further study and further development.

We note that the AutoDock Vina calculations, which are performed in vacuum, overpredict the presence of cooperative hydrophobic contacts and underestimate the populations of charge contacts and hydrogen bonds relative to explicit solvent MD. In contrast, the diffusion generative model of DiffDock appears to substantially underestimate the populations of all intermolecular interactions other than aromatic stacking. We observe that both AutoDock Vina and DiffDock generated more favorable docking scores for IDP conformations that adopted more compact, hairpin-like structures, with larger hydrophobic cores containing multiple aromatic residues. This suggests that both scoring functions could be further optimized or retrained to produce more accurate results for IDP ligands. Such efforts will benefit from the development of large experimental and computational benchmarks for predicting the affinity of IDP ligands to the binding sites of multiple IDPs. It will also be important to test and adapt ensemble docking approaches, like the ones proposed here, on IDPs where ligand binding has been found to more strongly modulate the apo ensemble (24, 49). In these systems, it may be necessary to develop flexible docking approaches that sample protein backbone and sidechain degrees of freedom in apo ensembles of IDPs to identify accurate binding poses.

In conclusion, the results of this study suggest that ensemble docking approaches show promise for predicting the affinities of small molecule drugs to IDPs and describing the heterogenous binding mechanisms of IDP ligands in atomic detail. IDP ensemble docking approaches therefore warrant further study and development and may ultimately provide a valuable high throughput tool for IDP drug discovery campaigns.

## Methods

### MD Simulations

Previously reported MD simulations of apo α-syn-C-term and α-syn-C-term in the presence of Fasudil, Ligand 47 and Ligand 23 were run at 300K in the NPT ensemble with the Anton supercomputer (11). α-syn-C-term, water and ions were parameterized using the a99SB-*disp* force field (41) and ligands with the generalized amber force field (GAFF1) (44). Bonds involving hydrogen atoms were restrained to their equilibrium bond length and nonbonded interactions were truncated at 10A. Simulations were performed in a cubic water box with a length of 42 Å per side with one copy of α-syn-C-term and one copy of a ligand, corresponding to protein and small-molecule concentrations of 0.020M. Na or Cl ions were added to a concentration of 25mM for Ligands 23 and 47, and to a concentration of 50mM for Fasudil. A simulation of apo α-syn-C-term was run for 100μs and simulations of α-syn-C-term in the presence of Fasudil and Ligand 47 were run for 200μs. A simulation of α-syn-C-term in the presence of Ligand 23 was run for 60μs.

### t-SNE clustering

Each long-time scale MD simulation was clustered using a recently described t-distributed stochastic neighbor embedding (t-SNE) clustering method (45). Each trajectory was clustered into 20 structurally unique clusters, using a grid search to identify the t-SNE perplexity value that produced the highest silhouette score as previously described (45). Perplexity values of 1200, 1100 and 1800 were used to cluster Fasudil, Ligand 47 and Ligand 23, respectively. In protein MD ensembles simulated with a ligand, conformations where no ligand atom was within 6Å of the protein were discarded following clustering. After removing unbound frames, 1000 conformations were randomly selected without replacement from each t-SNE cluster for docking, resulting in an ensemble of 20000 conformations for ensemble docking. In the case of Ligand 23, the ligand-bound α-syn-C-term ensemble used as input for holo docking contained only 18,861 ligand-bound conformations, as some t-SNE clusters obtained from MD contained fewer than 1000 ligand-bound conformations. For these clusters, we holo docking was performed on all available ligand-bound conformations.

### AutoDock Vina ensemble docking

In AutoDock Vina ensemble docking calculations, the ligand of interest is docked on every residue of the protein. Sidechains are kept rigid, and the ligand was allowed torsional degrees of freedom. The search space was centered on the center of mass of each residue in a volume proportional to the ligand’s radius of gyration in accordance with established AutoDock Vina protocols (46). The radii of gyration of Ligand 47, Fasudil, and Ligand 23 are 4.23 Å, 3.48 Å, and 3.53 Å, corresponding to AutoDock Vina search space volumes of 4561 Å^3^, 2220 Å^3^, and 2646 Å^3^, respectively. Docking at each α-syn-C-term residue resulted in 20 possible ligand bound conformations for each protein conformation. We selected the pose with the most favorable AutoDock Vina docking score as the final docked pose for each α-syn-C-term conformation.

### DiffDock Ensemble docking

In DiffDock ensemble docking calculations, the ligand was docked only once on each protein conformation using the default DiffDock parameters. We utilize the score of DiffDock’s trained confidence model as the docking score of each predicted pose.

### Analysis of IDP ligand binding modes

To evaluate the similarity of ligand binding modes in docked ensembles and long-time scale MD simulations, we employ several approaches to characterize protein-ligand interactions in MD ensembles and docked ensembles. We performed these analyses on each α-syn-C-term conformational state identified by t-SNE – and used the population of each t-SNE cluster from MD to compute ensemble averages for each docked ensemble.

We calculate the per-residue contact probability between ligands and each residue in α-syn-C-term, defining a contact as occurring in all frames where at least one heavy (non-hydrogen) ligand atom is within 6.0 Å of a heavy protein atom of a given residue. We also calculate a *dual-residue contact probability* for all pairs of residues, which describes the probability that a pair of residues simultaneously form ligand contacts in any frame. To quantify the similarity of binding modes in docked ensembles and MD ensembles, we calculate the Pearson correlation coefficient (*r*) and RMSE of the per-residue contact probabilities and dual-residue contact probabilities observed in each α-syn-C-term conformational state in docked ensembles and MD ensembles.

We compute the populations of intermolecular protein-ligand hydrophobic contacts, aromatic stacking interactions, charge-charge contacts, and hydrogen bonds formed with each residue of α-syn-C-term as previously defined (11). Briefly, we define hydrophobic contacts as occurring when any ligand and protein carbon atoms (excluding protein Cα atoms) are within 5Å. We define charge contacts as occurring when any two atoms with opposite formal charges are within 5Å. Hydrogen bonds were identified as any potential hydrogen bond donor (hydrogen attached to nitrogen, oxygen, or sulfur) within 3.5Å of a heavy-atom hydrogen bond acceptor, with a donor hydrogen-acceptor angle > 150°. We define aromatic stacking based on distance and geometric orientation of ligand aromatic groups and aromatic protein sidechain chains: we define a vector R connecting the centroids of the protein and ligand aromatic groups and define a stacking interaction as occurring when the length of R is less than 5Å and the angle between the vector R and vectors formed by the normal of the planes of the each aromatic group are both less than 45°.

### Docked-pose RMSD calculations

RMSD calculations of ligand atomic positions are a common metric used to compare docked ligand poses to experimental ligand positions (28, 29). To compare the similarity of ligand poses in docked ensembles and ligand poses sampled in MD ensembles, we calculated the RMSD of the atomic positions of heavy (non-hydrogen) ligand atoms using two approaches. In the first approach, which we refer to as a *frame-matched RMSD*, we directly compute the ligand RMSD between an MD bound pose and a docked pose for a each frame in a docked ensemble. In frame-matched RMSD calculations, the protein coordinates are identical in the docked pose and ligand pose. We align the structures on the protein coordinates and directly compute the ligand RMSD between poses.

In the second approach, which we refer to as the *best-matched RMSD* of a docked pose, we identify the ligand RMSD of the most similar ligand-bound pose sampled in an MD ensemble. For a docked pose in a t-SNE cluster, we align the Cα protein coordinates of all ligand-bound conformations present in the corresponding t-SNE cluster of an MD simulation and compute the ligand RMSD from each aligned MD frame. We identify the MD frame with the smallest ligand RMSD as the best-matched MD frame and report the ligand RMSD of this frame as the best-matched RMSD for the docked pose.

### Comparing docking scores

We compared relative docking scores to known experimental affinities of ligands (11) using a scale we define as a *normalized docking score*. AutoDock Vina and DiffDock report different ranges of docking scores with different meanings (28, 29). To account for this, and enable comparisons in differences in the relative docking scores obtained within each set of docking calculations (AutoDock Vina holo docking, AutoDock Vina apo docking, DiffDock holo docking, DiffDock apo docking), we normalize all docking scores obtained from a given method from 0 to 1 using min-max normalization:

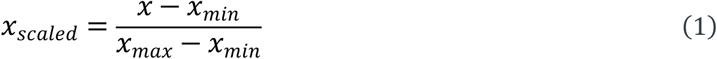

where *x_min_* and *x_max_* are worst and best docking scores observed in all docking calculations performed with Ligand 47, Fasudil and Ligand 23. With this normalization, the most favorable docking score in a set of docking calculations has a value of 1.0 and the least favorable docking score has a value of 0.

## Data & Code Availability

All code used for ensemble docking and trajectory analyses and all docked ensembles are freely available from GitHub (https://github.com/paulrobustelli/Dhar_IDP_ensemble_docking_25/). The α-syn-C-term MD trajectories analyzed here are available for non-commericial use by request from D.E. Shaw Research.

## Acknowledgements

This work was supported by the National Institutes of Health under award R35GM142750. A.D. addtionally acknowledges the support of a John L. Zabriskie Jr. ‘61 Undergraduate Research Fellowship. T.R.S additionally acknowledges the support of a GAANN Fellowship from the Department of Education (GAANN P200A240037). We thank Dr. Jaya Krishna Koneru for providing support with t-SNE clustering calculations.

