## Supporting Information for "Ensemble docking for intrinsically disordered proteins"

### Supplementary Information

Anjali Dhar, Thomas R. Sisk & Paul Robustelli[†]

Dartmouth College, Department of Chemistry, Hanover, NH, 03755

† To whom correspondence should be addressed.

Paul Robustelli:

Address: 6128 Burke Laboratory
Department of Chemistry
Hanover, NH, 03755

| **Average normalized docking score (uncertainty)** | **Autodock Vina Holo Docking** | **DiffDock Holo Docking** | **AutoDock Vina Apo Docking** | **DiffDock Apo Docking** |
|---|---|---|---|---|
| Ligand 47 | 0.326 (0.0016) | 0.630 (0.0011) | 0.289 (0.0013) | 0.672 (0.0011) |
| Fasudil | 0.249 (0.0016) | 0.594 (0.0012) | 0.237 (0.0015) | 0.639 (0.0013) |
| Ligand 23 | 0.220 (0.0015) | 0.554 (0.0017) | 0.228 (0.0014) | 0.588 (0.0016) |

**Supplementary Table 1. Average normalized docking scores reported by AutoDock Vina and DiffDock for all ensemble docking approaches.** For each set of docking calculations (holo docking or apo docking) with each docking program (AutoDock Vina or DiffDock) docking scores of all ligands are min-max normalized onto a single normalized docking score scale using the highest and lowest docking scores observed in calculations of all three ligands. Normalized docking scores are therefore not directly comparable between AutoDock Vina and DiffDock calculations or between apo and holo docking calculations with each method. We report the ensemble-average value of the normalized docking scores in each docked ensemble and report an uncertainty estimate of the average normalized docking score obtained from bootstrapping using 10000 samples for each docked ensemble. Uncertainty estimates were determined by averaging the magnitude of the upper and lower deviations of the 95% confidence interval in each sample, and we calculate the mean of this value over all bootstrapping samples.

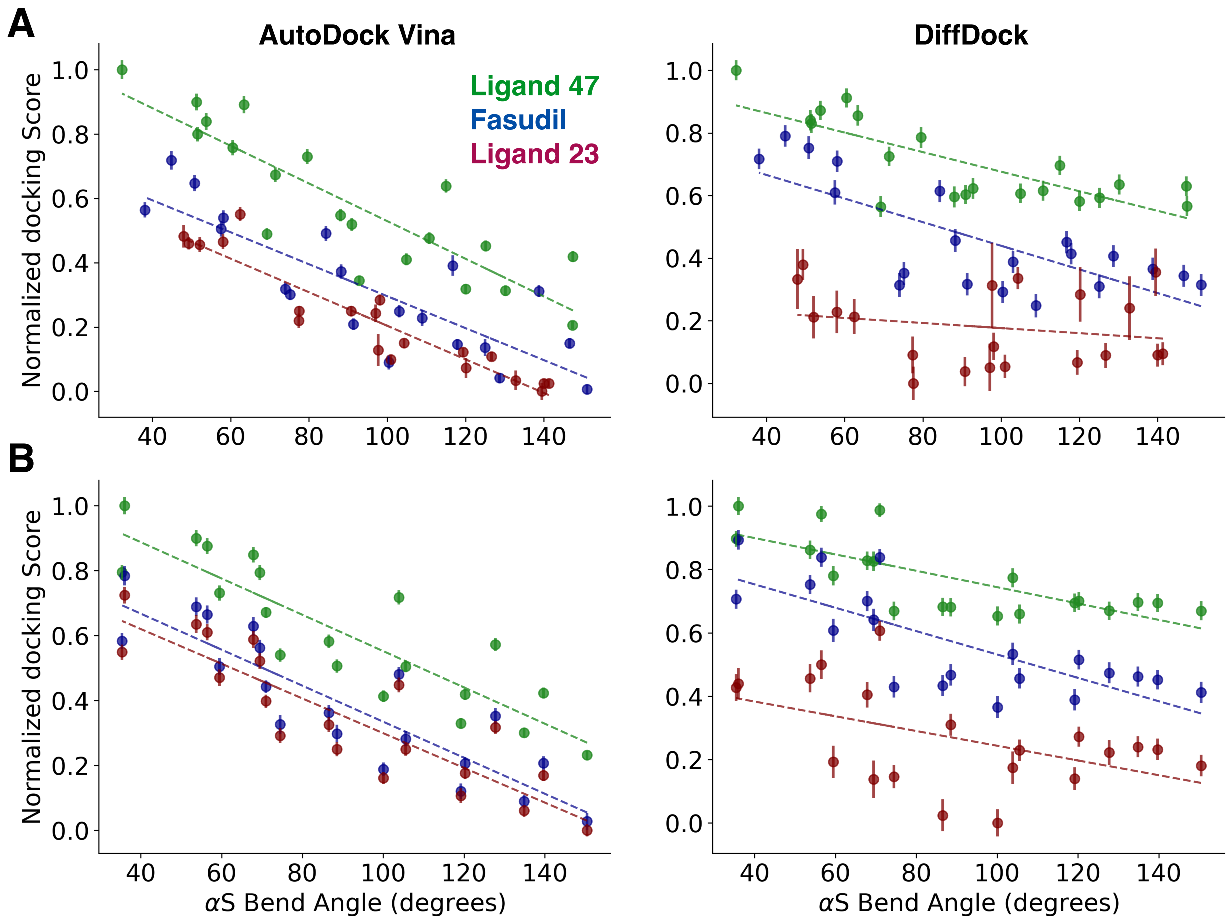


**Supplementary Figure 1. Relationship between the average α-syn-C-term bend angle and docking scores in holo docked ensembles (A) and apo docked ensembles (B) obtained from AutoDock Vina and DiffDock ensemble docking.** Docking scores for all three ligands were min-max normalized for each set of docking calculations, and the average normalized docking score was calculated for each cluster identified by t-SNE. The α-syn-C-term bend angle (αS bend angle) was defined as the angle formed by the Cα atoms of residues 121, 131 and 140 of α-syn-C-term. Error estimates are shown as vertical lines. Error estimates indicate the mean of the upper and lower deviations of the 95% confidence interval calculated from bootstrapping using 10000 samples.

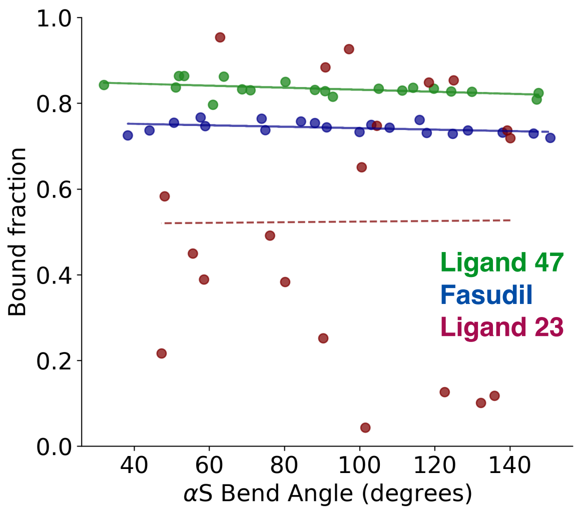


**Supplementary Figure 2. Relationship between the average α-syn-C-term bend angle and fraction of bound frames in long time scale MD simulation of α-syn-C-term with Ligand 47, Fasudil and Ligand 23.** Bound fractions of each t-SNE cluster were calculated for each t-SNE cluster identified from each MD simulation, where a bound pose is defined as any frame where a ligand heavy atom is within 6Å of any protein heavy atom. αS bend angle is defined as the angle formed by the Cα atoms of residues 121, 131 and 140 of α-syn-C-term.

| $r^2$ | Autodock Vina Holo Docking | DiffDock Holo Docking | AutoDock Vina Apo Docking | DiffDock Apo Docking | Long timescale MD |
|---|---|---|---|---|---|
| Ligand 47 | 0.75 | 0.62 | 0.77 | 0.57 | 0.22 |
| Fasudil | 0.70 | 0.56 | 0.78 | 0.59 | 0.18 |
| Ligand 23 | 0.88 | 0.04 | 0.78 | 0.25 | 0.00 |

**Supplementary Table 2. Goodness of fit ($r^2$) values of the linear relationship between the average α-syn-C-term bend angle and the average normalized docking of each t-SNE cluster obtained with each ensemble-docking approach.** The goodness of fit of the linear relationship between the α-syn-C-term bend angle of each t-SNE cluster and the bound fraction observed in each cluster in long timescale MD simulations of each ligand are shown for comparison.

| *r* (RMSE) | | Ligand contacts | Hydrophobic contacts | Aromatic stacking | Hydrogen bonds | Charge contacts | Dual-residue contacts |
|---|---|---|---|---|---|---|---|
| Ligand 47 | *AutoDock Vina Holo* | 0.86 (0.26) | 0.90 (0.11) | 0.89 (0.05) | 0.40 (0.04) | 0.77 (0.02) | 0.90 (0.19) |
| | *AutoDock Vina Apo* | 0.70 (0.28) | 0.83 (0.11) | 0.60 (0.04) | 0.35 (0.04) | 0.61 (0.03) | 0.77 (0.20) |
| | *DiffDock Holo* | 0.71 (0.11) | 0.88 (0.07) | 0.05 (0.05) | -0.09 (0.03) | 0.39 (0.04) | 0.80 (0.06) |
| | *DiffDock Apo* | 0.68 (0.12) | 0.87 (0.07) | 0.16 (0.06) | -0.07 (0.03) | 0.40 (0.04) | 0.75 (0.07) |
| Fasudil | *AutoDock Vina Holo* | 0.81 (0.27) | 0.87 (0.12) | 0.42 (0.09) | 0.25 (0.02) | 0.61 (0.02) | 0.87 (0.18) |
| | *AutoDock Vina Apo* | 0.70 (0.28) | 0.80 (0.13) | 0.14 (0.09) | 0.24 (0.02) | 0.54 (0.02) | 0.79 (0.19) |
| | *DiffDock Holo* | 0.50 (0.13) | 0.80 (0.06) | -0.16 (0.06) | -0.17 (0.03) | 0.31 (0.03) | 0.74 (0.06) |
| | *DiffDock Apo* | 0.51 (0.13) | 0.81 (0.06) | -0.11 (0.06) | -0.16 (0.03) | 0.57 (0.03) | 0.72 (0.06) |
| Ligand 23 | *AutoDock Vina Holo* | 0.32 (0.34) | 0.44 (0.16) | 0.00 (0.07) | 0.34 (0.06) | | 0.52 (0.22) |
| | *AutoDock Vina Apo* | 0.30 (0.36) | 0.35 (0.29) | 0.27 (0.06) | 0.31 (0.03) | | 0.47 (0.23) |
| | *DiffDock Holo* | 0.16 (0.22) | 0.41 (0.09) | 0.88 (0.05) | 0.22 (0.01) | | 0.43 (0.10) |
| | *DiffDock Apo* | 0.19 (0.23) | 0.38 (0.10) | 0.66 (0.05) | 0.22 (0.1) | | 0.42 (0.10) |
| Average | *AutoDock Vina Holo* | 0.66 (0.29) | 0.74 (0.13) | 0.42 (0.07) | 0.33 (0.04) | 0.68 (0.02) | 0.76 (0.20) |
| | *AutoDock Vina Apo* | 0.57 (0.31) | 0.65 (0.18) | 0.32 (0.06) | 0.29 (0.03) | 0.57 (0.03) | 0.65 (0.21) |
| | *DiffDock Holo* | 0.46 (0.15) | 0.69 (0.08) | 0.27 (0.05) | -0.02 (0.02) | 0.37 (0.04) | 0.66 (0.07) |
| | *DiffDock Apo* | 0.46 (0.16) | 0.70 (0.08) | 0.25 (0.06) | 0.01 (0.02) | 0.49 (0.04) | 0.63 (0.08) |

**Supplementary Table 3**. **Pearson correlation coefficients and root-mean square error (RMSE) values of per-residue populations of intermolecular contacts observed in docked ensembles and ligand bound ensembles obtained from long-time scale MD simulations of α-syn-C-term.** We compute the Pearson correlation coefficients and RMSE values of the per-residue populations of intermolecular interactions observed in docked ensembles and ligand-bound MD ensembles in each of the 20 clusters identified by t-SNE and report the average values across all clusters. RMSE values are displayed in parentheses. Ligand 23 contains no formal charges – so no charge contacts are present.

| *r* (RMSE) | | Ligand contacts | Hydrophobic contacts | Aromatic stacking | Hydrogen bonds | Charge contacts | Dual-residue contacts |
|---|---|---|---|---|---|---|---|
| Ligand 47 | *AutoDock Vina Holo* | 0.91 (0.24) | 0.94 (0.10) | 0.97 (0.03) | 0.51 (0.03) | 0.91 (0.02) | 0.94 (0.17) |
| | *AutoDock Vina Apo* | 0.90 (0.23) | 0.89 (0.09) | 0.49 (0.04) | 0.39 (0.03) | 0.68 (0.02) | 0.90 (0.16) |
| | *DiffDock Holo* | 0.80 (0.08) | 0.92 (0.06) | -0.56 (0.02) | -0.06 (0.03) | 0.62 (0.03) | 0.86 (0.04) |
| | *DiffDock Apo* | 0.83 (0.08) | 0.89 (0.06) | 0.44 (0.03) | -0.07 (0.03) | 0.46 (0.04) | 0.84 (0.05) |
| Fasudil | *AutoDock Vina Holo* | 0.84 (0.25) | 0.91 (0.11) | 0.71 (0.08) | 0.27 (0.02) | 0.74 (0.02) | 0.91 (0.17) |
| | *AutoDock Vina Apo* | 0.87 (0.25) | 0.76 (0.11) | -0.78 (0.09) | 0.14 (0.02) | 0.47 (0.02) | 0.85 (0.16) |
| | *DiffDock Holo* | 0.58 (0.11) | 0.88 (0.05) | -0.39 (0.04) | -0.20 (0.03) | 0.41 (0.03) | 0.80 (0.05) |
| | *DiffDock Apo* | 0.59 (0.11) | 0.76 (0.07) | -0.96 (0.06) | -0.23 (0.03) | 0.24 (0.03) | 0.70 (0.05) |
| Ligand 23 | *AutoDock Vina Holo* | 0.83 (0.30) | 0.93 (0.13) | 0.94 (0.12) | 0.78 (0.06) | | 0.88 (0.20) |
| | *AutoDock Vina Apo* | 0.88 (0.30) | 0.70 (0.26) | -0.65 (0.09) | 0.84 (0.03) | | 0.87 (0.19) |
| | *DiffDock Holo* | 0.60 (0.15) | 0.84 (0.05) | -0.71 (0.05) | 0.76 (0.00) | | 0.75 (0.07) |
| | *DiffDock Apo* | 0.68 (0.14) | 0.80 (0.06) | -0.94 (0.06) | 0.74 (0.01) | | 0.73 (0.07) |
| Average | *AutoDock Vina Holo* | 0.86 (0.26) | 0.93 (0.11) | 0.87 (0.08) | 0.52 (0.04) | 0.83 (0.02) | 0.91 (0.18) |
| | *AutoDock Vina Apo* | 0.88 (0.26) | 0.78 (0.15) | -0.31 (0.07) | 0.46 (0.03) | 0.58 (0.02) | 0.87 (0.17) |
| | *DiffDock Holo* | 0.66 (0.11) | 0.88 (0.05) | -0.55 (0.04) | 0.17 (0.02) | 0.52 (0.03) | 0.80 (0.05) |
| | *DiffDock Apo* | 0.70 (0.11) | 0.82 (0.06) | -0.49 (0.05) | 0.15 (0.02) | 0.35 (0.04) | 0.76 (0.06) |

**Supplementary Table 4. Pearson correlation coefficients and root-mean square error (RMSE) values of ensemble-averaged per-residue populations of intermolecular contacts observed in docked ensembles and ligand bound ensembles obtained from long-time scale MD simulations of α-syn-C-term.** We compute the Pearson correlation coefficients and RMSE values of the ensemble-averaged per-residue populations of intermolecular interactions observed in docked ensembles and ligand-bound MD ensembles. RMSE values are displayed in parentheses.

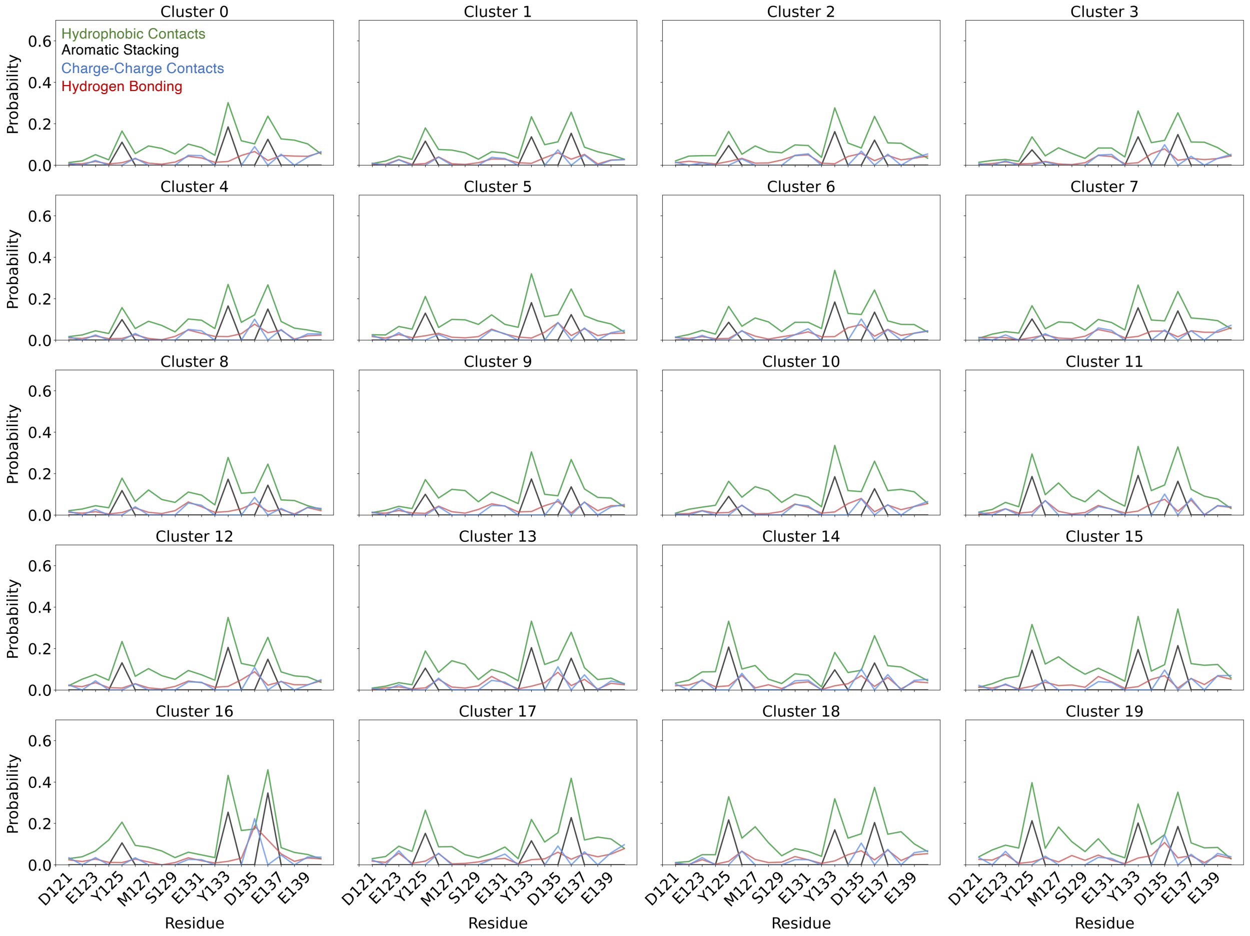


**Supplementary Figure 3. Per-residue populations of intermolecular interactions between Ligand 47 and α-syn-C-term in each t-SNE cluster from a 200μs MD simulation of α-syn-C-term with Ligand 47.** Hydrophobic contacts are displayed in green, aromatic stacking interactions are displayed in black, hydrogen bonds are displayed in red, and charge contacts are displayed in blue.

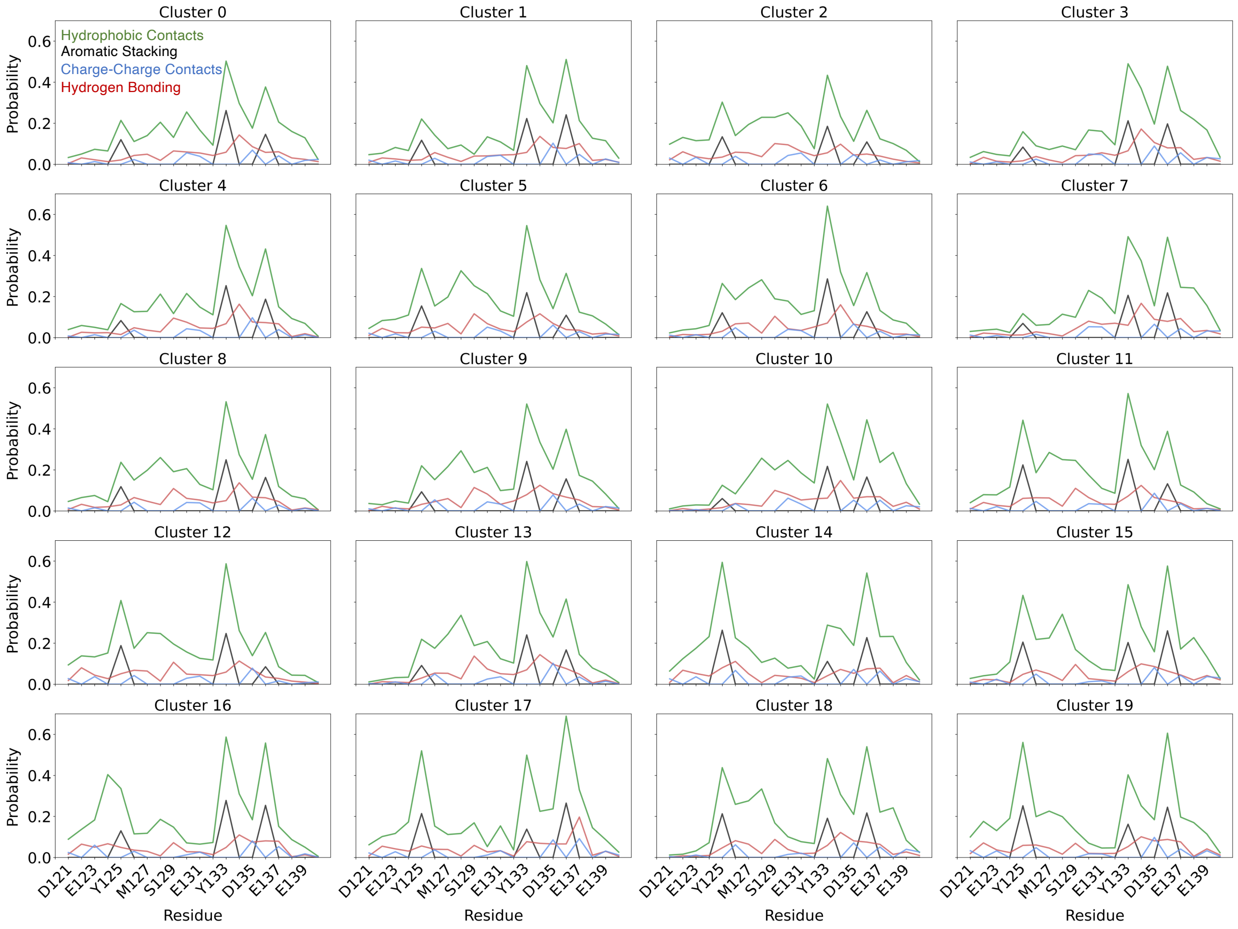


**Supplementary Figure 4. Per-residue populations of intermolecular interactions between Ligand 47 and α-syn-C-term obtained from AutoDock Vina holo docking on each t-SNE cluster of α-syn-C-term identified from a 200μs MD simulation of α-syn-C-term with Ligand 47.** Hydrophobic contacts are displayed in green, aromatic stacking interactions are displayed in black, hydrogen bonds are displayed in red, and charge contacts are displayed in blue.

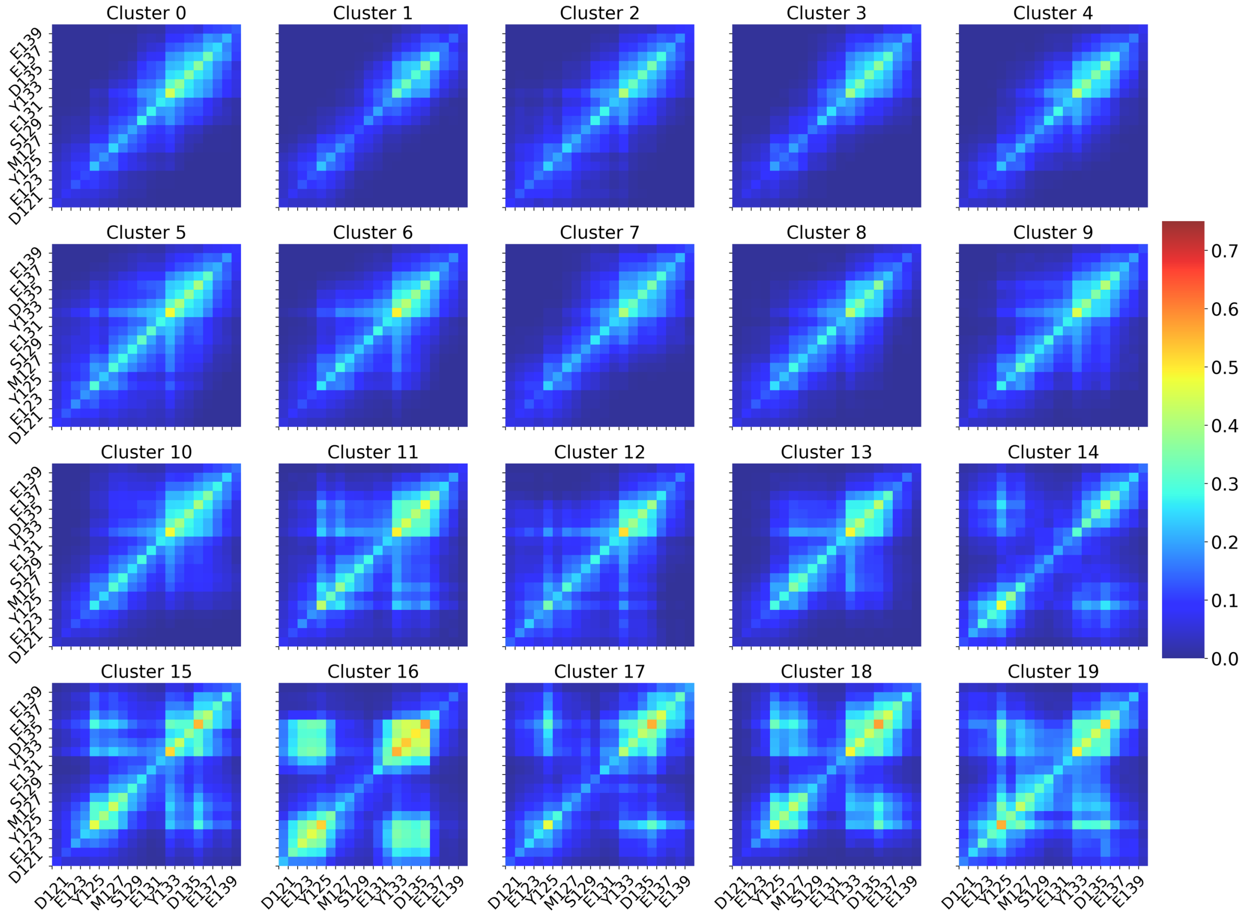


**Supplementary Figure 5. Populations of dual-residue contact probability between Ligand 47 and α-syn-C-term in each t-SNE cluster from a 200μs MD simulation of α-syn-C-term with Ligand 47.** The dual residue contact condition is met when any ligand heavy atom is within 6.0 Å of any two heavy atoms in a pair of residues in α-syn-C-term simultaneously.

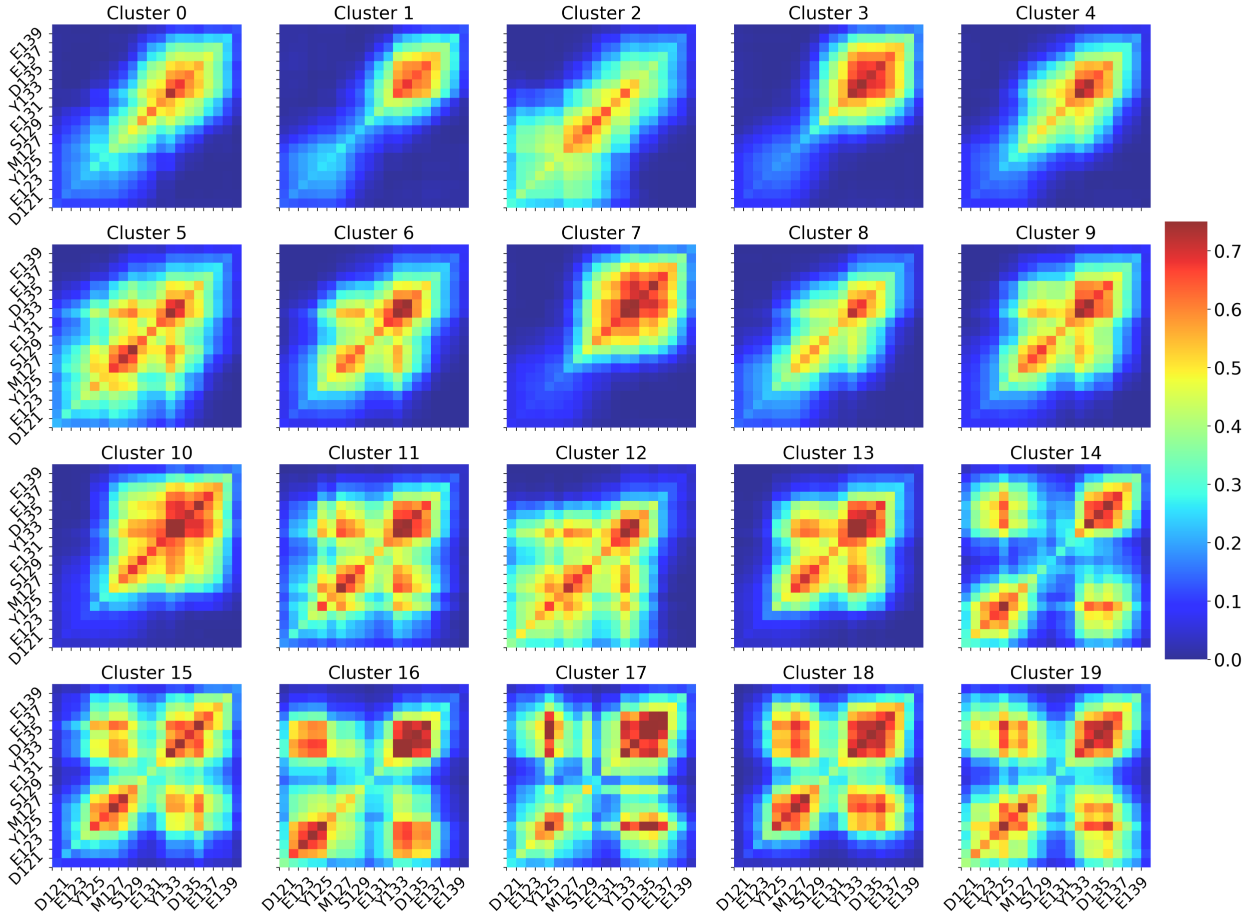


**Supplementary Figure 6. Populations of dual-residue contact probability between Ligand 47 and α-syn-C-term obtained from AutoDock Vina holo docking on each t-SNE cluster of α-syn-C-term identified from a 200μs MD simulation of α-syn-C-term with Ligand 47.** The dual residue contact condition is met when any ligand heavy atom is within 6.0 Å of any two heavy atoms in a pair of residues in α-syn-C-term simultaneously.

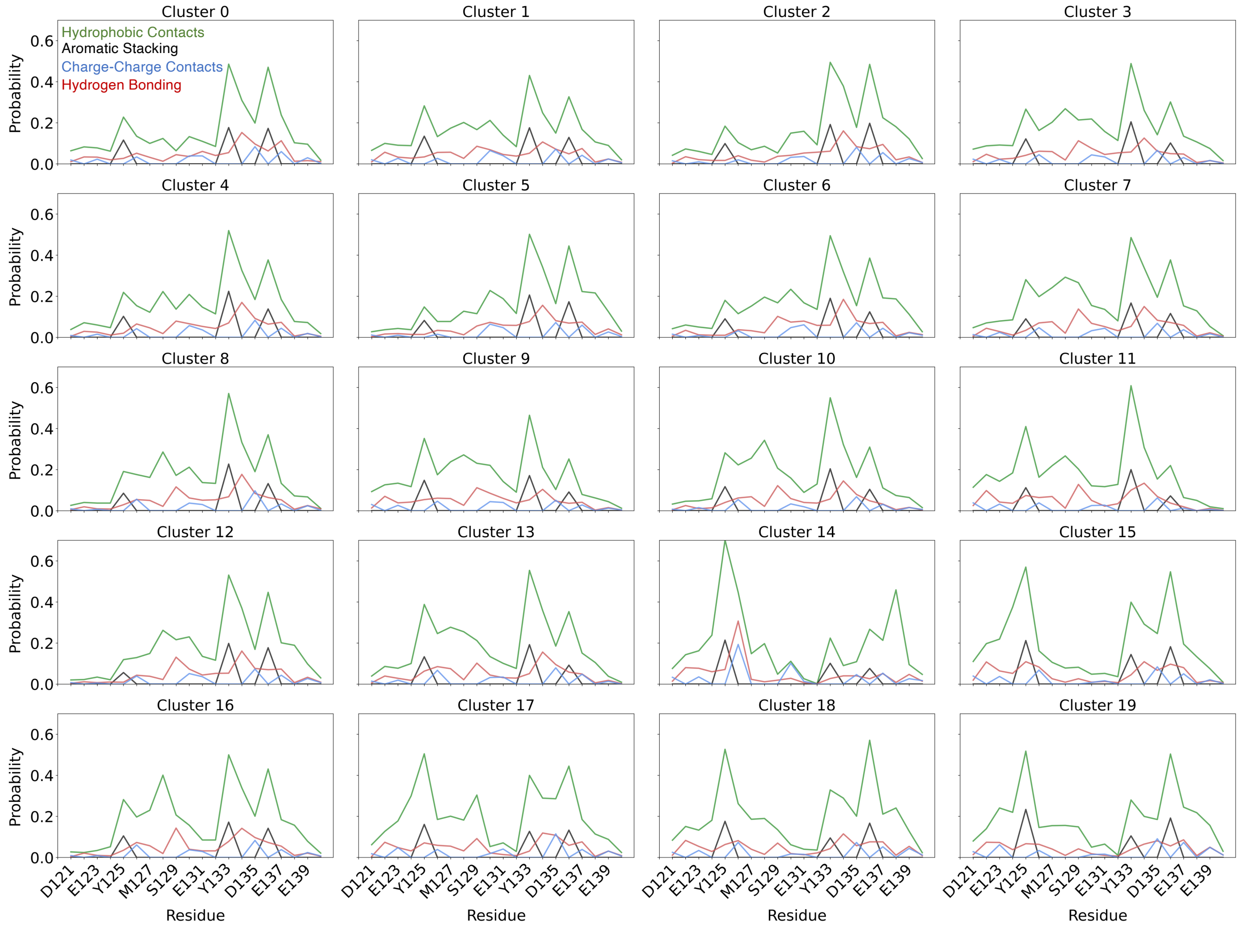


**Supplementary Figure 7. Per-residue populations of intermolecular interactions between Ligand 47 and α-syn-C-term obtained from AutoDock Vina apo docking on each t-SNE cluster of α-syn-C-term identified from a 100μs MD simulation of α-syn-C-term.** Hydrophobic contacts are displayed in green, aromatic stacking interactions are displayed in black, hydrogen bonds are displayed in red, and charge contacts are displayed in blue.

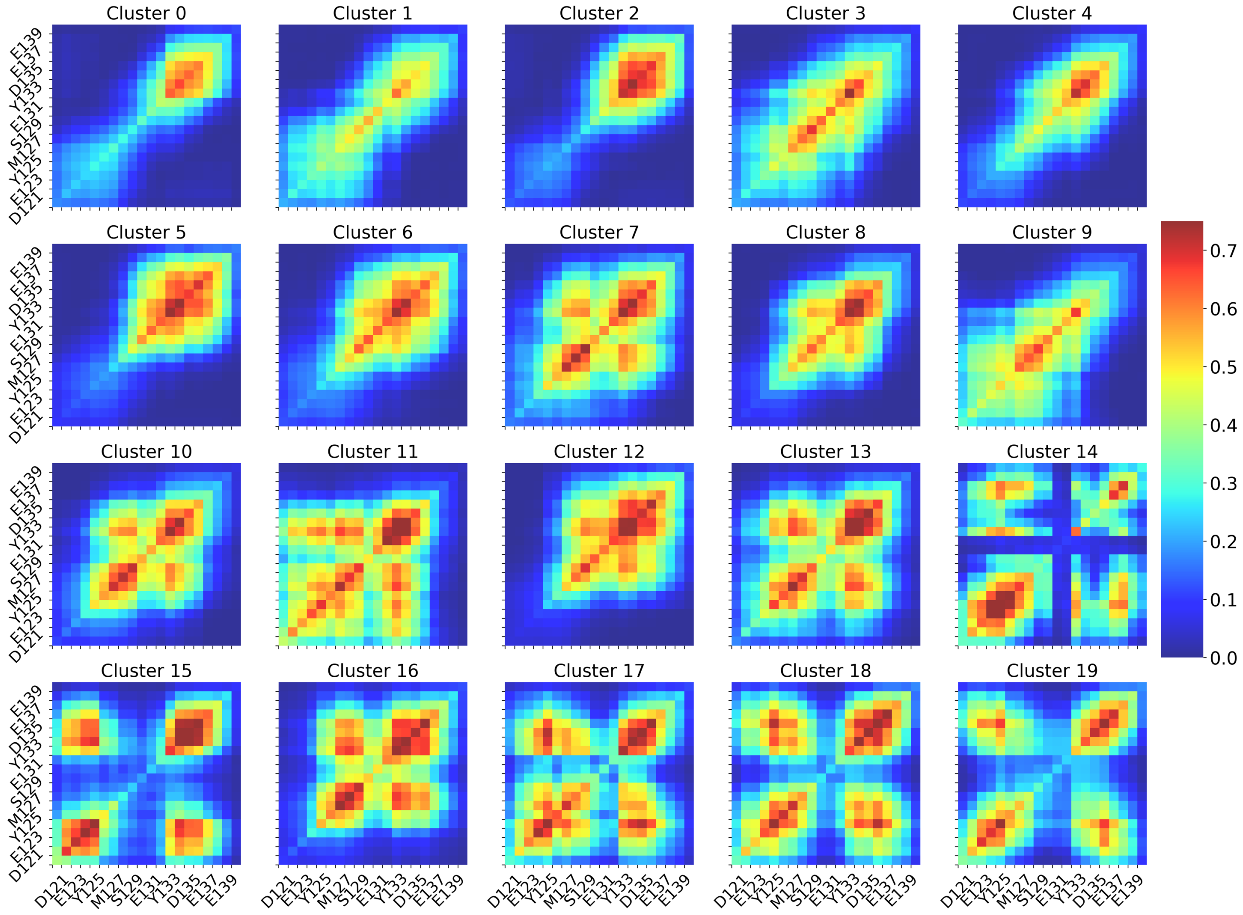


**Supplementary Figure 8. Populations of dual-residue contact probability between Ligand 47 and α-syn-C-term obtained from AutoDock Vina apo docking on each t-SNE cluster of α-syn-C-term identified from a 100μs MD simulation of α-syn-C-term.** The dual residue contact condition is met when any ligand heavy atom is within 6.0 Å of any two heavy atoms in a pair of residues in α-syn-C-term simultaneously.

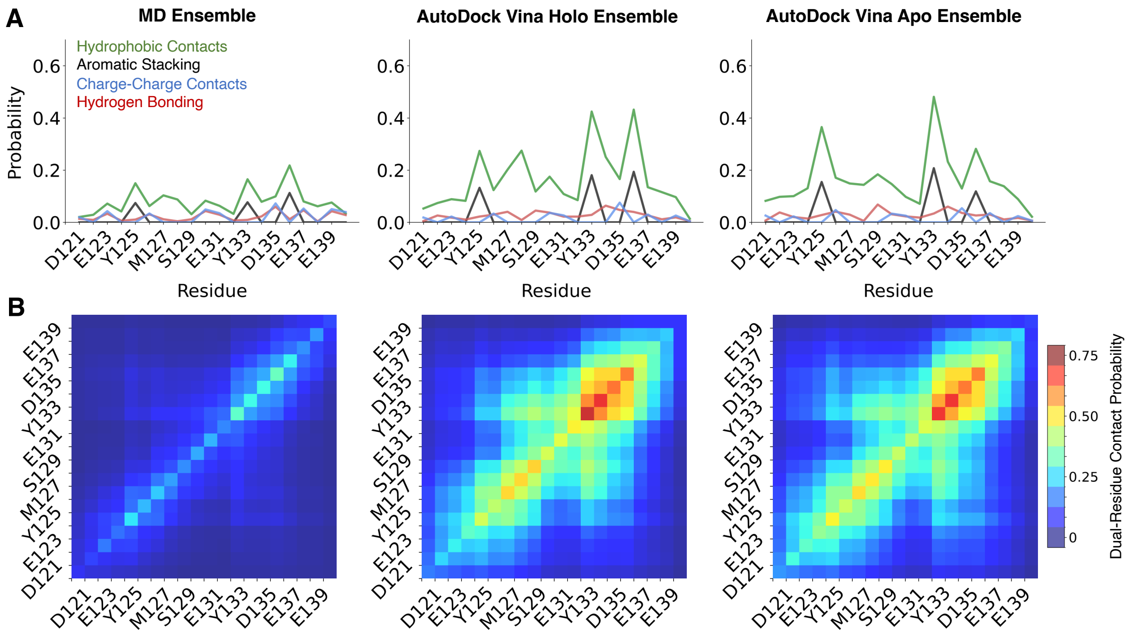


**Supplementary Figure 9. Comparison of ensemble-averaged intermolecular protein-ligand interactions (A) and dual-residue contact probabilities (B) between Fasudil and α-syn-C-term obtained from a 200μs MD simulation with Fasudil, AutoDock Vina holo docking, and AutoDock Vina apo docking.**

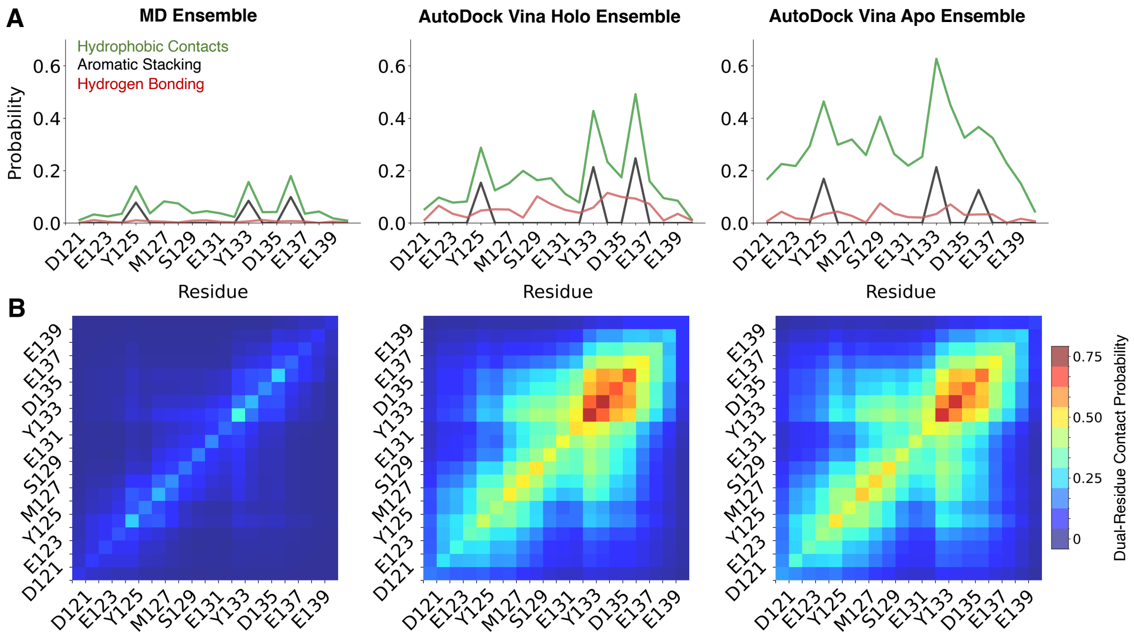


**Supplementary Figure 10. Comparison of ensemble-averaged intermolecular protein-ligand interactions (A) and dual-residue contact probabilities (B) between Ligand 23 and α-syn-C-term obtained from a 60μs MD simulation with Ligand 23, AutoDock Vina holo docking, and AutoDock Vina apo docking.**

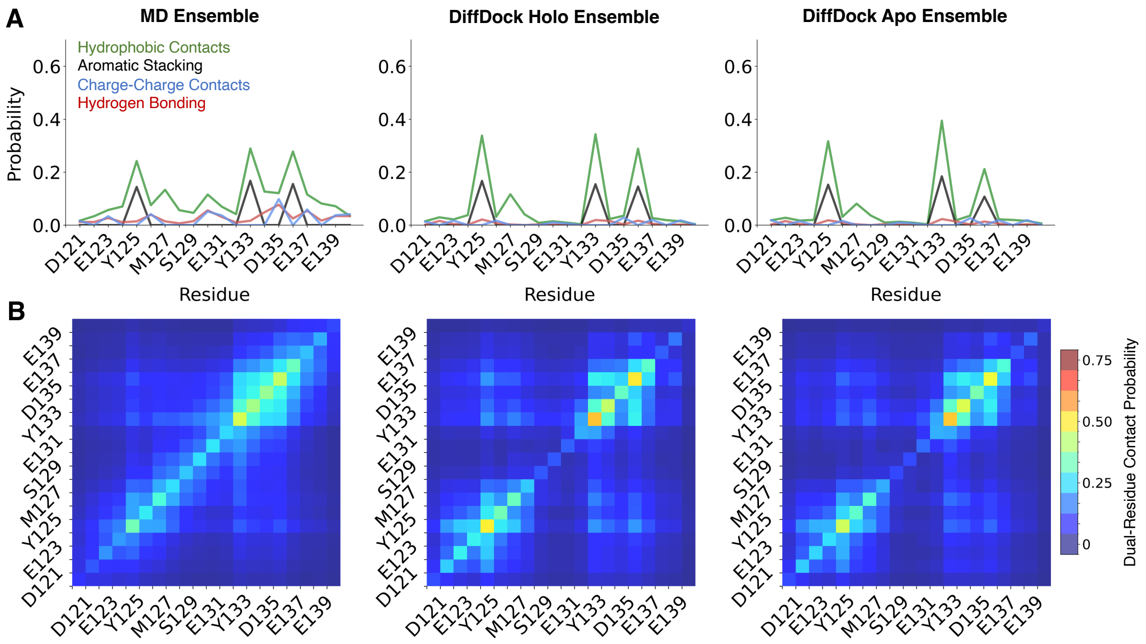


**Supplementary Figure 11. Comparison of ensemble-averaged intermolecular protein-ligand interactions (A) and dual-residue contact probabilities (B) between Ligand 47 and α-syn-C-term obtained from a 200μs MD simulation with Ligand 47, DiffDock holo docking, and DiffDock apo docking.**

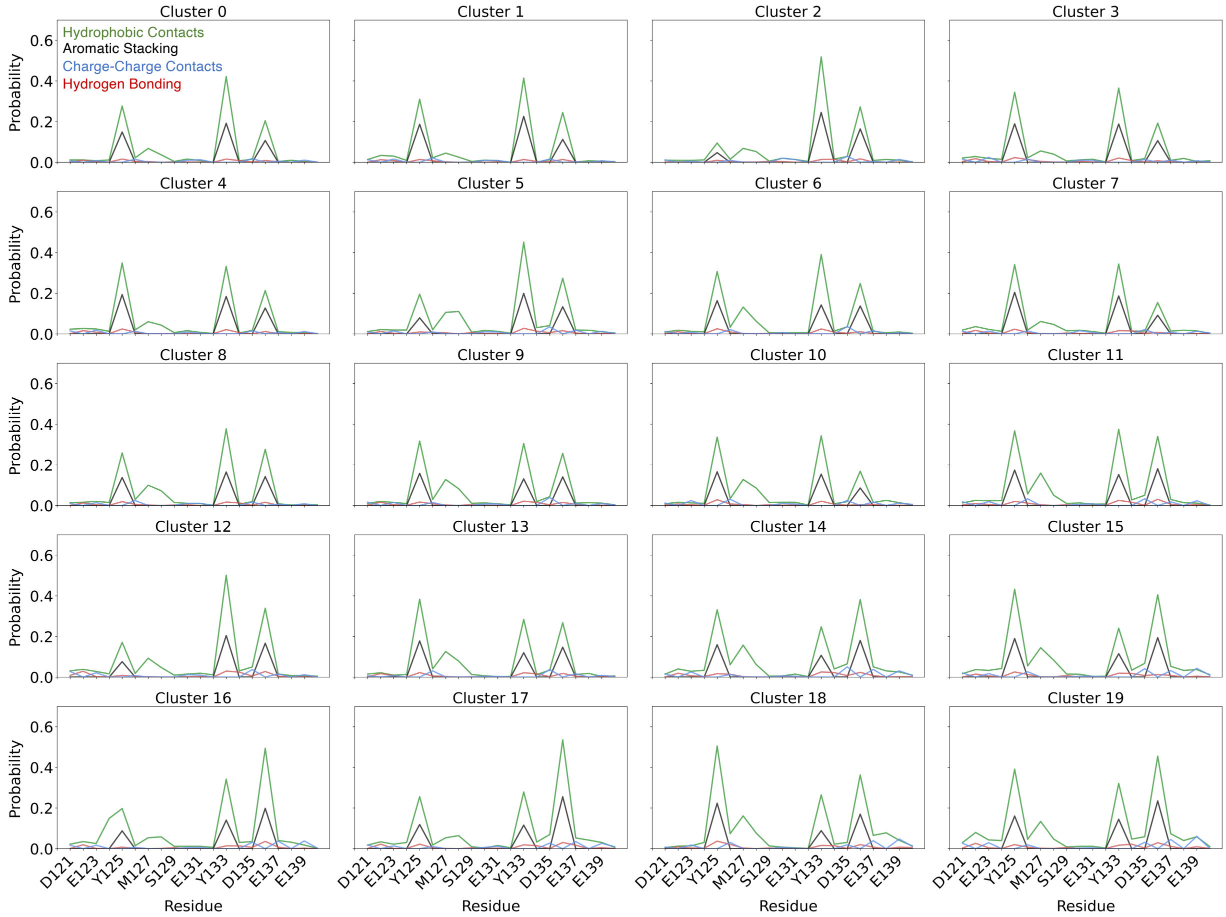


**Supplementary Figure 12. Per-residue populations of intermolecular interactions between Ligand 47 and α-syn-C-term obtained from DiffDock holo docking on each t-SNE cluster of α-syn-C-term identified from a 200μs MD simulation of α-syn-C-term with Ligand 47.** Hydrophobic contacts are displayed in green, aromatic stacking interactions are displayed in black, hydrogen bonds are displayed in red, and charge contacts are displayed in blue.

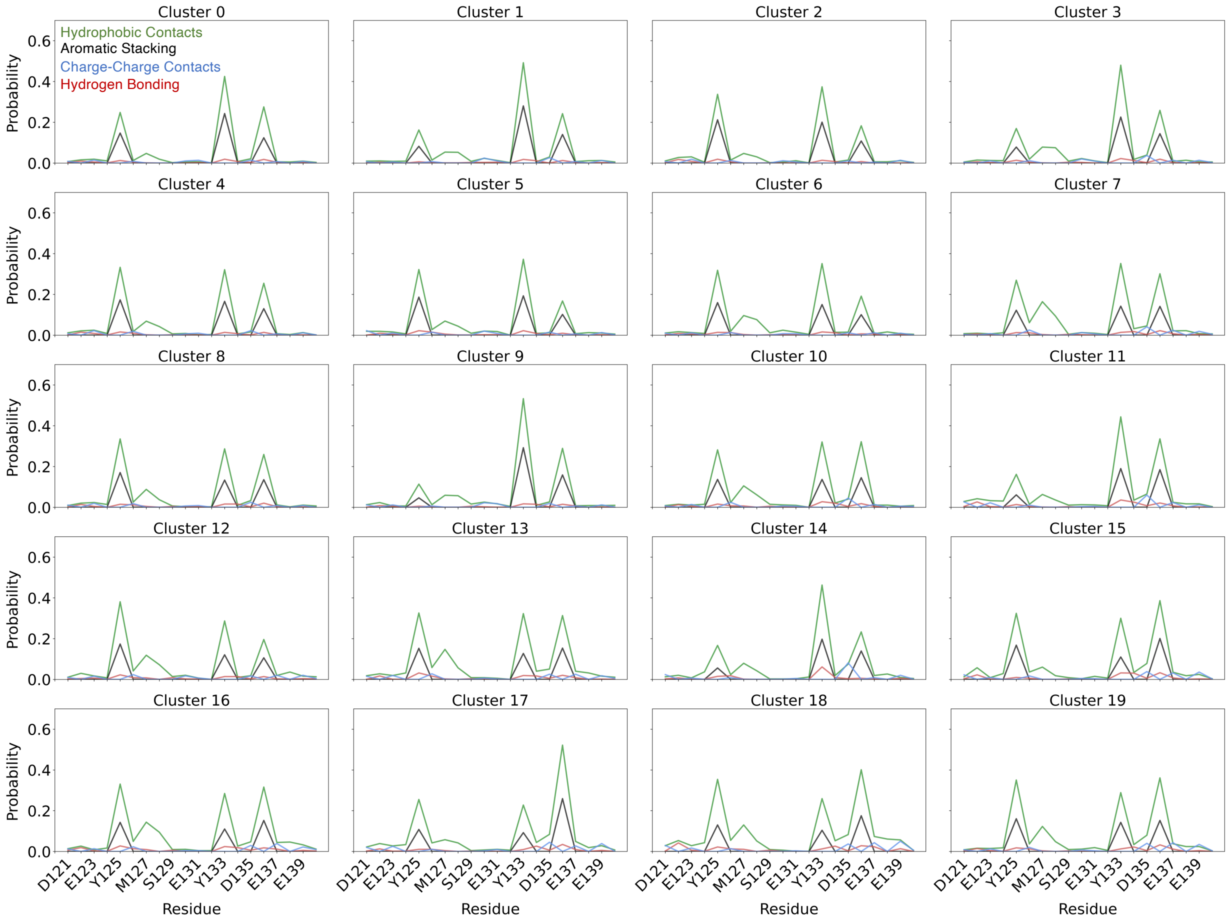


**Supplementary Figure 13. Per-residue populations of intermolecular interactions between Ligand 47 and α-syn-C-term obtained from DiffDock apo docking on each t-SNE cluster of α-syn-C-term identified from a 100μs MD simulation of α-syn-C-term.** Hydrophobic contacts are displayed in green, aromatic stacking interactions are displayed in black, hydrogen bonds are displayed in red, and charge contacts are displayed in blue.

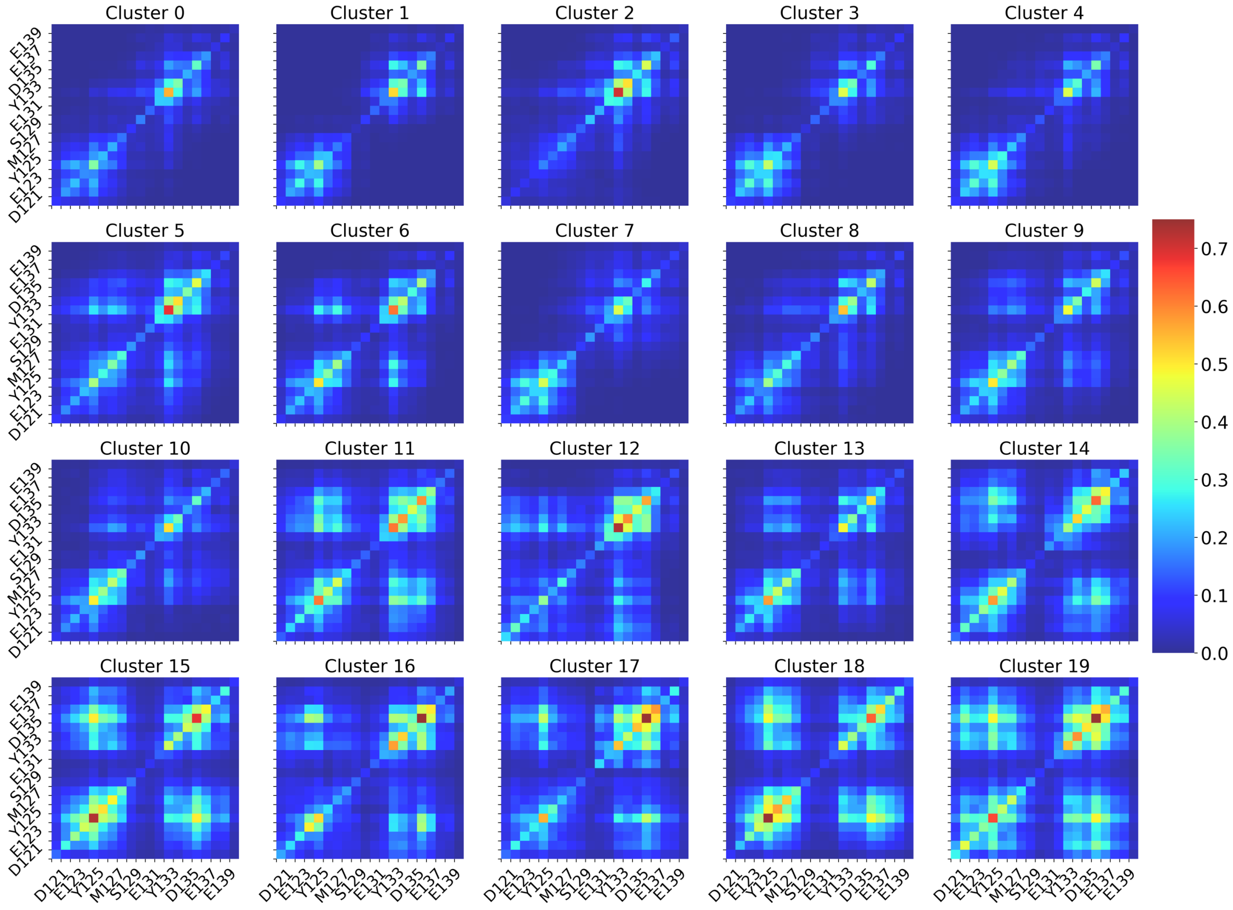


**Supplementary Figure 14. Populations of dual-residue contact probability between Ligand 47 and α-syn-C-term obtained from DiffDock holo docking on each t-SNE cluster of α-syn-C-term identified from a 200μs MD simulation of α-syn-C-term with Ligand 47.** The dual residue contact condition is met when any ligand heavy atom is within 6.0 Å of any two heavy atoms in a pair of residues in α-syn-C-term simultaneously.

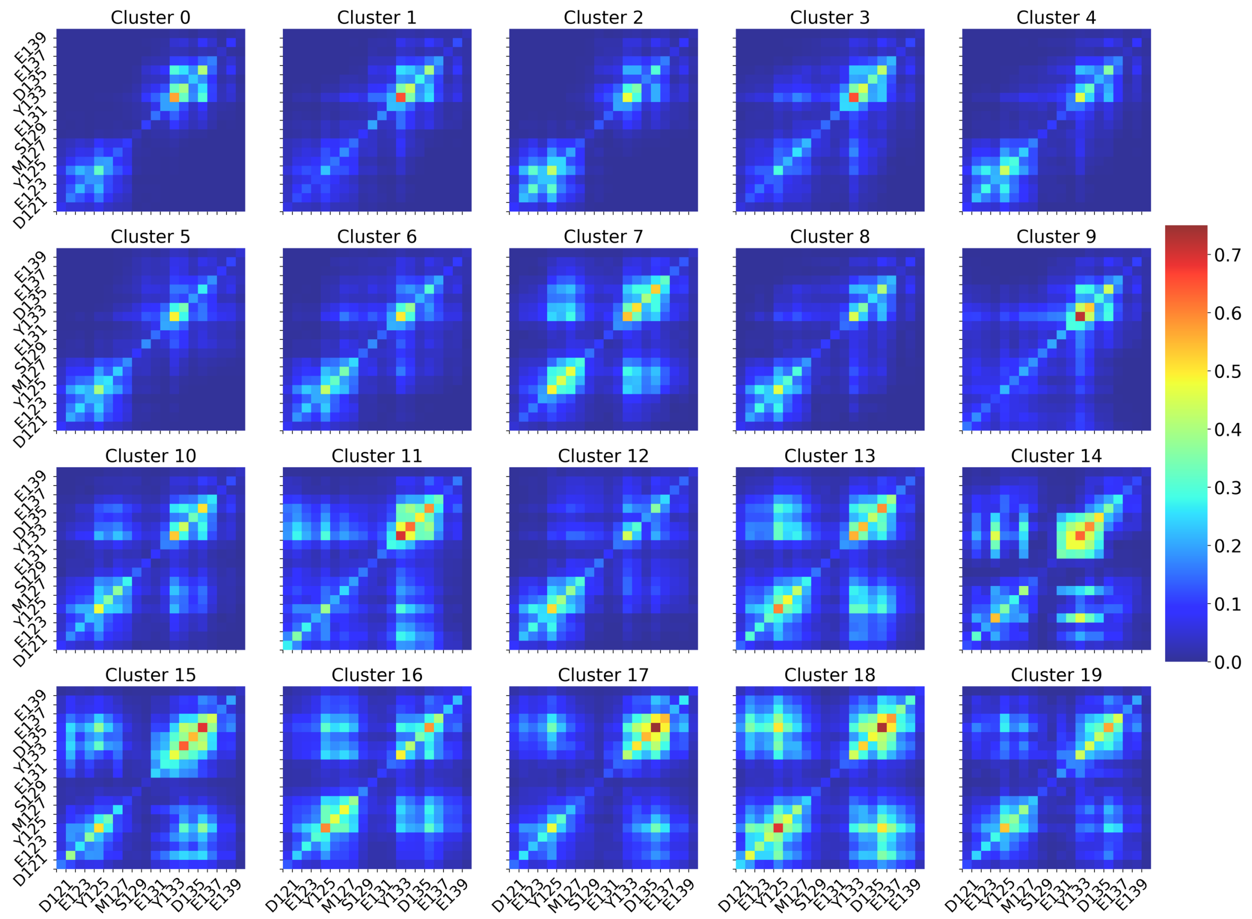


**Supplementary Figure 15. Populations of dual-residue contact probability between Ligand 47 and α-syn-C-term obtained from DiffDock apo docking on each t-SNE cluster of α-syn-C-term identified from a 100μs MD simulation of α-syn-C-term.** The dual residue contact condition is met when any ligand heavy atom is within 6.0 Å of any two heavy atoms in a pair of residues in α-syn-C-term simultaneously.

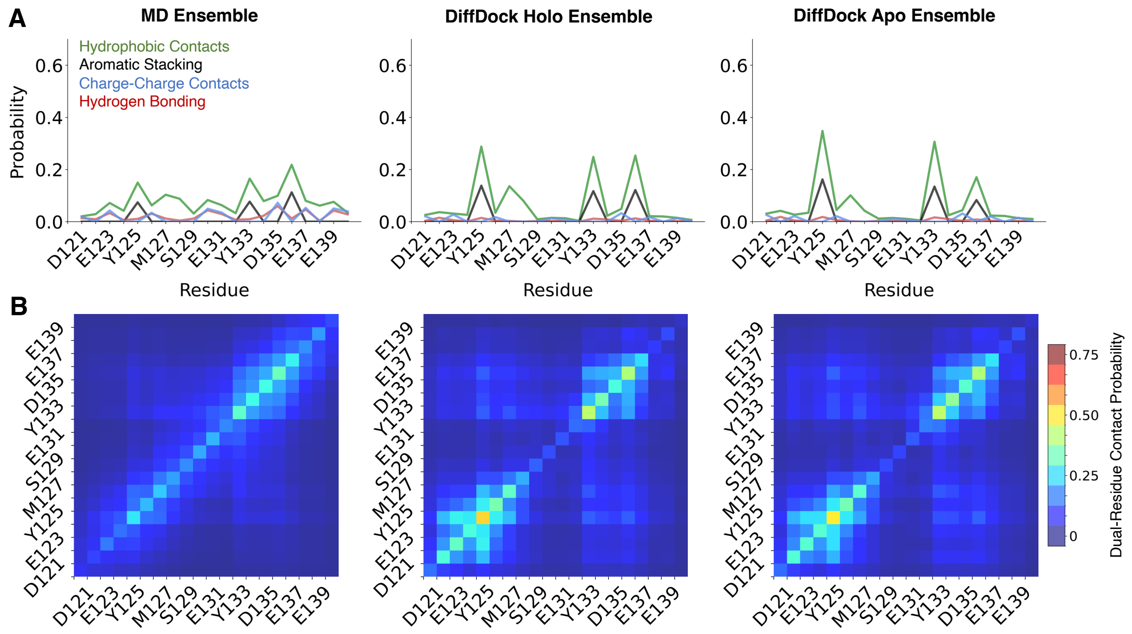


**Supplementary Figure 16. Comparison of ensemble-averaged intermolecular protein-ligand interactions (A) and dual-residue contact probabilities (B) between Fasudil and α-syn-C-term obtained from a 200μs MD simulation with Fasudil, DiffDock holo docking, and DiffDock apo docking.**

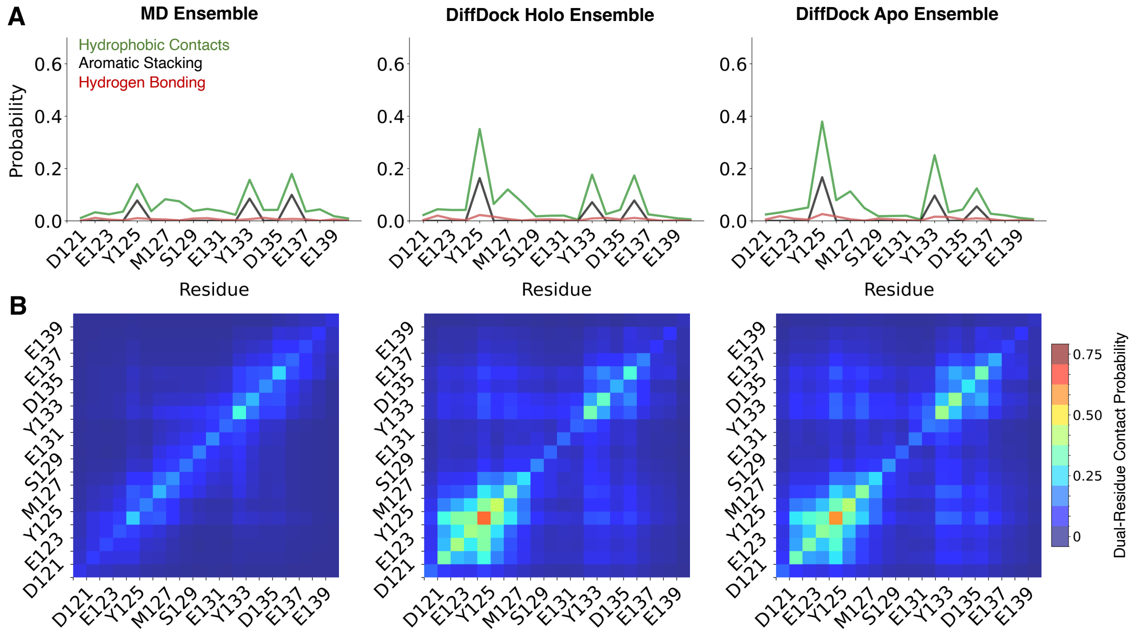


**Supplementary Figure 17. Comparison of ensemble-averaged intermolecular protein-ligand interactions (A) and dual-residue contact probabilities (B) between Ligand 23 and α-syn-C-term obtained from a 60μs MD simulation with Ligand 23, DiffDock holo docking, and DiffDock apo docking.**

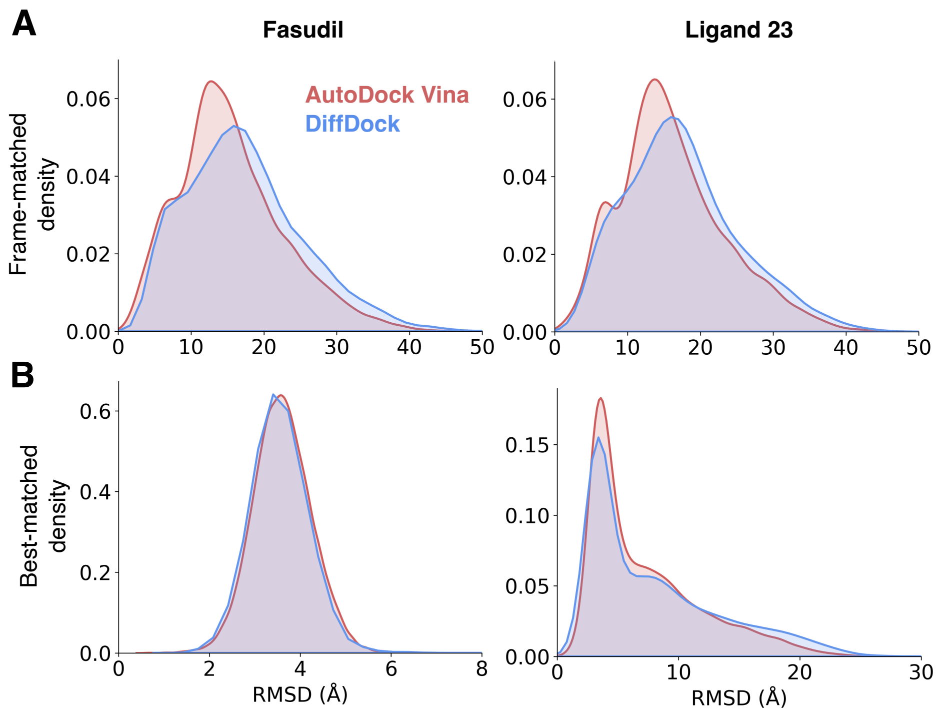


**Supplementary Figure 18. Distributions of ligand RMSD of ligand poses predicted by docking and references poses observed in long time scale MD simulation for Fasudil and Ligand 23 holo docking with the frame-matched analysis method (A) and the best-matched analysis method (B).** Predicted docking poses are obtained from holo docking with either AutoDock Vina (red) or DiffDock (blue), where each ligand is docked on the long-time scale MD simulation of α-syn-C-term that was originally simulated with the same ligand.

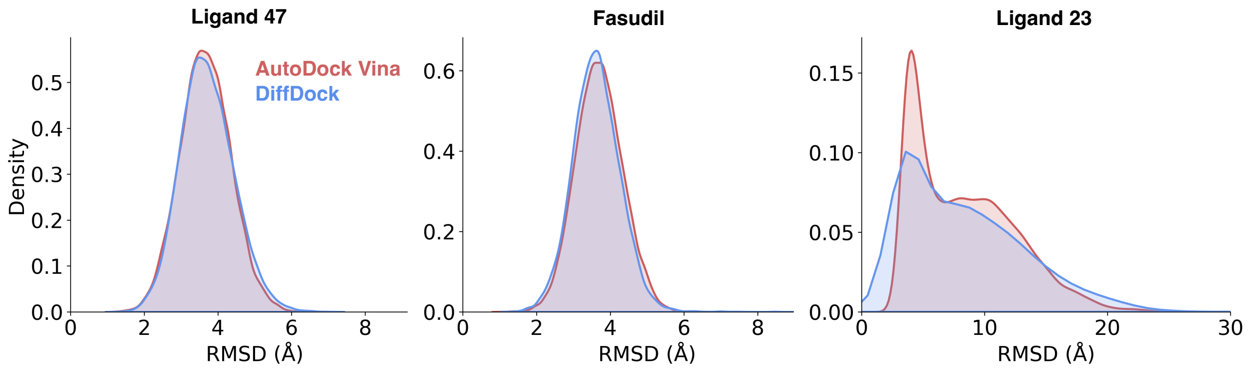


**Supplementary Figure 19. Distributions of best-matched ligand RMSD of ligand poses predicted by apo docking and references poses observed in long time scale MD simulation for Ligand 47, Fasudil and Ligand 23.** Predicted docking poses are obtained from apo docking with either AutoDock Vina (red) or DiffDock (blue), where each ligand was docked on the 100μs MD simulation of α-syn-C-term that was originally simulated with no ligand. Reference ligand poses were obtained from the long-time scale MD simulation of α-syn-C-term that were originally simulated with the docked ligand.

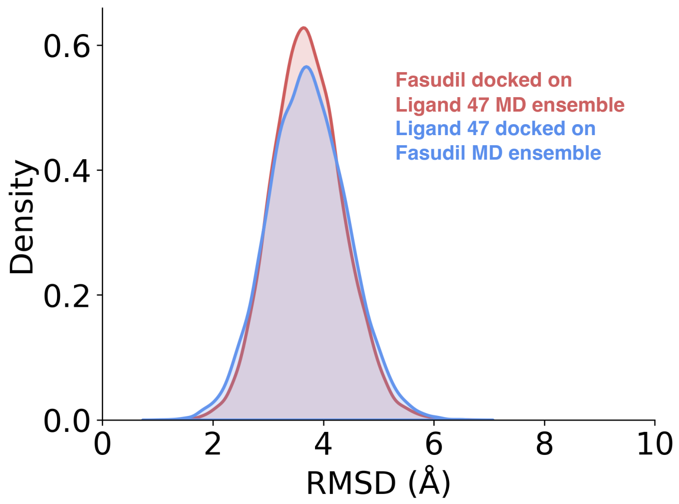


**Supplementary Figure 20. Distribution of best-matched ligand RMSD of ligand poses predicted by docking ligands on α-syn-C-term conformations obtained from a holo MD ensemble obtained from a different ligand.** Cross docking is defined as docking a ligand on a long-time scale MD simulation of α-syn-C-term with a different ligand than was originally simulated. Reference ligand poses for cross docked ensembles were obtained from the long-time scale MD simulation of α-syn-C-term that were originally simulated with the docked ligand.

| Percentage of frames with RMSD < 3 Å (RMSD < 5Å) | Autodock Vina Docked on Fasudil Ensemble | Autodock Vina Docked on Ligand 47 Ensemble |
|---|---|---|
| Ligand 47 | 15.36 (96.13) | 27.96 (98.30) |
| Fasudil | 16.54 (98.42) | 13.11 (97.34) |

**Supplementary Table 5. Best-matched ligand RMSD of ligand poses predicted by docking and reference poses observed in long-time scale MD simulation for cross docked ensembles.** Cross docking is defined as docking a ligand on a long-time scale MD simulation of α-syn-C-term with a different ligand than was originally simulated. We report the percentage of docked frames obtained by holo or cross docking where the best-matched RMSD of ligand heavy-atom coordinates is less than 3Å from the reference MD bound-pose and less than 5Å from the reference MD bound-pose (in parentheses). References for cross docking runs were obtained from the long-time scale MD simulation of α-syn-C-term with the docked ligand.

| Average normalized docking score (uncertainty) | Autodock Vina Docked on Fasudil Ensemble | Autodock Vina Docked on Ligand 47 Ensemble |
|---|---|---|
| Ligand 47 | 0.235 (0.0014) | 0.329 (0.0016) |
| Fasudil | 0.248 (0.0016) | 0.223 (0.0016) |

**Supplementary Table 6. Average normalized docking scores reported by AutoDock Vina for cross docking approaches.** Cross docking is defined as docking a ligand on a long-time scale MD simulation of α-syn-C-term with a different ligand than was originally simulated. For each set of docking calculations (holo docking or cross docking), docking scores of Ligand 47 and Fasudil docked on the same MD ensemble of α-syn-C-term are min-max normalized onto a single normalized docking score using the highest and lowest docking scores observed. Normalized docking scores are only directly comparable between ligands docked on the same MD ensemble of α-syn-C-term. We report the ensemble-average value of the normalized docking scores in each docked ensemble and report an uncertainty estimate of the average normalized docking score obtained from bootstrapping using 10000 samples for each docked ensemble. Uncertainty estimates were determined by averaging the magnitude of the upper and lower deviations of the 95% confidence interval in each sample, and we calculate the mean of this value over all bootstrapping samples.
